# DLK-dependent axonal mitochondrial fission drives degeneration following axotomy

**DOI:** 10.1101/2023.01.30.526132

**Authors:** Jorge Gómez-Deza, Matthew Nebiyou, Mor R. Alkaslasi, Francisco M. Nadal-Nicolás, Preethi Somasundaran, Anastasia L. Slavutsky, Michael E. Ward, Wei Li, Trent A. Watkins, Claire E. Le Pichon

## Abstract

Currently there are no effective treatments for an array of neurodegenerative disorders to a large part because cell-based models fail to recapitulate disease. Here we developed a reproducible human iPSC-based model where laser axotomy causes retrograde axon degeneration leading to neuronal cell death. Time-lapse confocal imaging revealed that damage triggers an apoptotic wave of mitochondrial fission proceeding from the site of injury to the soma. We demonstrated that this apoptotic wave is locally initiated in the axon by dual leucine zipper kinase (DLK). We found that mitochondrial fission and resultant cell death are entirely dependent on phosphorylation of dynamin related protein 1 (DRP1) downstream of DLK, revealing a new mechanism by which DLK can drive apoptosis. Importantly, we show that CRISPR mediated *Drp1* depletion protected mouse retinal ganglion neurons from degeneration after optic nerve crush. Our results provide a powerful platform for studying degeneration of human neurons, pinpoint key early events in damage related neural death and new focus for therapeutic intervention.

## Introduction

Neuronal death and axon loss are universal hallmarks of neurodegeneration, relevant to Alzheimer’s disease, ALS and other neurodegenerative diseases. However, our understanding of the mechanisms controlling axon degeneration and neuron death is incomplete. Advances in this area have enormous potential to benefit all types of neurodegenerative conditions including those arising from nerve injury and traumatic brain injury.

Separate mechanisms regulate axon degeneration and neuronal apoptosis [1]. For example, deletion of BAX (BCL-2-associated X protein), a central regulator of apoptosis, does not protect axons from Wallerian degeneration. On the other hand, BAX deletion protects neurons following trophic factor withdrawal, a developmental model of axon degeneration and death [2–4] . SARM1 is the central executioner of Wallerian axon degeneration [5, 6]. However, it is unclear how universal this pathway is for axon degeneration in the context of disease since some neurodegenerative models are not rescued by loss of SARM1 [7, 8]. Furthermore, while most neurodegenerative diseases involve loss of axons and cell bodies, Wallerian axon degeneration occurs independently from neuronal cell death.

Mitochondria are a central hub of axon health regulation, at the nexus of axonal energy production, oxidative stress, and calcium buffering [9, 10]. They also play critical roles in degeneration and regeneration [11, 12]. Healthy maintenance of the mitochondrial network is mediated by a balance of mitochondrial fission and fusion regulated by specialized machinery [13]. Interestingly, mutations in some of these genes cause neurodegenerative conditions such as motor and sensory neuropathy (eg. Mfn1 in Charcot-Marie-Tooth [14]). A key player in the mitochondrial fission process is dynamin related protein 1 or DRP1, a GTPase that oligomerizes around the outer mitochondrial membrane to cause mitochondrial fission [15]. DRP1 also regulates apoptosis [13, 16–18]. Under conditions that drive apoptosis, DRP1 promotes a type of apoptotic mitochondrial fission in which BAX permeabilizes the outer mitochondrial membrane, allowing the release of pro-apoptotic factors such as cytochrome c [19, 20].

Much of what we know about mechanisms regulating neuron degeneration and death comes from a wealth of studies of apoptosis in cell lines (but not neurons), or from animal models (neurons, but not human). How translatable this work is and how applicable to neurons remains to be determined. Recent advances in iPSC-derived neuron techniques now enable the generation of a diverse range of human neuron subtypes. These efforts have already generated exciting findings in the neurodegeneration field [21–23]. We therefore set out to study axon injury using a widely adopted glutamatergic neuron model produced by directed differentiation of human iPSCs (i3Neurons [24, 25]).

We and others have shown that the dual leucine zipper kinase (DLK, or MAP3K12), is a key regulator of axon degeneration and a driver of neuron cell death [26–30]. The canonical DLK pathway involves a phosphorylation cascade downstream of DLK homodimerization and a transcriptional stress response, including phosphorylation of the transcription factor cJun [30].

Despite its consideration as a drug target for several neurological conditions [31–33], relatively little is known about DLK function in human neurons.

Here, we investigate the behavior of i3Neurons following laser axotomy, and the role of DLK in the injury response of human neurons. Following axotomy, we observe progressive retrograde axon degeneration from the injury site back towards the cell body, causing subsequent neuronal death, and establishing a unique human model of neurodegeneration. This contrasts with the behavior of cultured mouse neurons, which survive and regenerate after this type of injury [34, 35], but resembles the pattern of neurodegeneration observed in optic nerve injuries in living mice [36] .

We find that in axons proximal to the injury, mitochondria shrink and undergo DRP1-dependent fission in a retrograde wave from the damage site, leading to apoptosis of the cell body. DLK regulates mitochondrial fission via phosphorylation of DRP1 at serine 616, defining a new substrate of the DLK signaling pathway. Such DRP1 phosphorylation is necessary for localization of BAX to axonal mitochondria, which causes axon degeneration. Blocking DLK or DRP1 protects axons from degeneration and neurons from cell death both in human i3Neurons *in vitro* and *in vivo* in the mouse. This study identifies a novel signaling mechanism for DLK that drives local axon degeneration without the need for transcription. Our findings reveal an apoptotic cascade that is initiated in the axon that both underlies a form of non-Wallerian axon degeneration and drives neuronal cell death.

## Results

### Axotomy in human neurons causes both proximal and distal axon degeneration

Axotomy has often been used as a method to study axon degeneration. After axotomy, the distal portion of the axon undergoes Wallerian degeneration, a phenomenon controlled by the NADase SARM1, the key executioner of this type of axon degeneration [5, 6]. To study the response of human neurons to axotomy, we performed laser axotomies in i3Neurons that were sparsely transduced with a fluorescent protein to visualize individual neurons (Fig. 1a). The distal portion of these axons underwent Wallerian degeneration as expected (Fig. 1b-c). To our surprise, we observed that the portion of the axon proximal to the injury and still connected to the cell body underwent progressive retrograde axon degeneration (Fig. 1b-c). Because SARM1-dependent degeneration relies on the loss of the axon survival factor NMNAT2, which occurs when the axon is severed from the cell body, we hypothesized that the proximal axon degeneration we observed might not rely on SARM1 and could provide an opportunity to elucidate novel pathways of axon degeneration.

**Figure 1.**
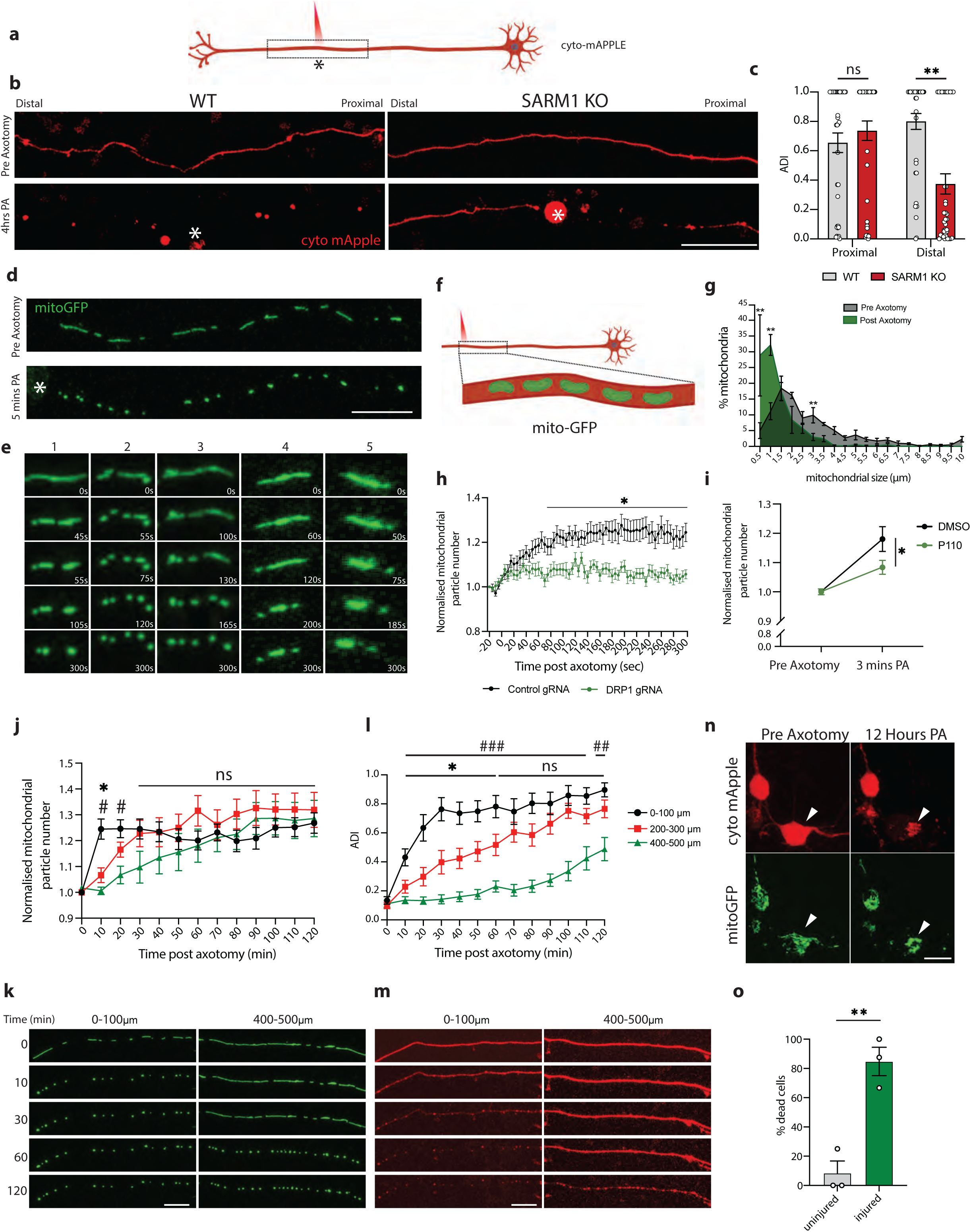
Axotomy triggers a wave of DRP1-dependent mitochondrial fission, axon degeneration and cell death. a) Schematic representation of i3Neuron laser axotomy using Biorender. Dashed box indicates field of view. * Marks the site of axotomy b) Representative images of WT and SARM1 KO axons transduced with cytoplasmic mApple (red) pre and 4 hours post axotomy (PA). * Marks the site of axotomy. (Scale bar = 100 µm). c) Quantification of axon degeneration index (ADI) in WT and SARM1 KO neurons distal and proximal to the site of axon injury 4 hours post axotomy. Results are represented as mean ± SEM. N=3 independent differentiations, N≥ 38 axons. Two-way ANOVA, Bonferroni correction (p≤0.01 **, not significant, ns). d) Representative images of i3Neuron axons transduced with mitoGFP (green) pre- and 5 minutes post axotomy (PA). * Marks the site of axotomy. (Scale bar = 20 µm). Time-lapse images of mitochondria shown in E. e) Representative images of mitochondria undergoing fission after axon injury outlined in D. f) Schematic representation of i3Neuron laser axotomy. Dashed box indicates field of view. g) Quantification of mitochondrial length pre (green) and 5 minutes post axotomy (grey) (PA). Results are represented as mean ± SEM. N=3 independent differentiations, N≥ 28 axotomized neurons. Two-way ANOVA, Bonferroni correction (p≤0.05 *). h) Normalized number of mitochondrial particles post axotomy in control and DRP1 KD neurons. Results are represented as mean ± SEM. N=3 independent differentiations, N≥ 25 axotomized neurons. Two-way ANOVA, Bonferroni correction (p≤0.05 *). i) Normalized number of mitochondrial particles 3 mins post axotomy in DMSO and P110 treated neurons. Results are represented as mean ± SEM. N=3 independent differentiations, N≥ 26 axotomized neurons. Two-way ANOVA, Bonferroni correction (p≤0.05 *). j) Quantification of axotomy-induced mitochondrial fission wave. Normalized number of mitochondrial particles post axotomy 0-100, 200-300 and 400-500 µm from the site of axotomy. Results are represented as mean ± SEM. N=3 independent differentiations, N≥ 23 axotomized neurons. One-way ANOVA, Bonferroni correction (*p≤0.05, 0-100 µm vs 200-300 µm, # p≤0.05, 0-100 µm vs 400-500 µm). k) Representative images of axotomy-induced mitochondrial fission wave, Pre axotomy (0), 10-, 30- and 120-minutes post axotomy. Neurons sparsely transduced with mitoGFP (green). (Scale bar = 25 µm). l) Quantification of axotomy-induced progressive axon degeneration. Axon degeneration index (ADI) 0-100, 200-300 and 400-500µm from the site of axotomy. Results are represented as mean ± SEM. N=3 independent differentiations, N≥ 14 axotomized neurons. (*p≤0.05, 0-100 µm vs 200-300 µm, ^##^ p≤0.01, ^###^ p≤0.005, 0-100 µm vs 400-500 µm). m) Representative images of axotomy-induced axon degeneration. Pre axotomy (0), 10-, 30- and 120-minutes post axotomy. Neurons sparsely transduced with cyto-mApple (red). * Marks the site of axotomy. (Scale bar = 25 µm). n) Representative images of axotomy-induced neuron death. Pre and 12-hours post axotomy. Arrow indicates axotomized neuron. Neurons sparsely transduced with cyto-mApple (red) and mitoGFP (green). (Scale bar = 40 µm). o) Quantification of axotomy-induced neuron death. Percentage of dead neurons 12 hours post axotomy. Results are represented as mean ± SEM. N=3 independent differentiations, N≥30 axotomized neurons. Unpaired t-test (p≤0.01 **).

We tested the requirement for SARM1 in the degeneration of axons both proximal and distal to the injury site. Using CRISPR/Cas9, we knocked out SARM1 in i3Neurons [37]. We then axotomized these neurons and control neurons and imaged the axon on both sides of the injury. As expected, SARM1 was necessary for distal, but not proximal, axon degeneration (Fig. 1b-c). We therefore investigated the mechanisms underlying proximal axon degeneration in these neurons. For the remainder of the study, we focus specifically on the axon segment proximal to the injury site.

### Axotomy in human neurons triggers a wave of DRP1-dependent mitochondrial fission

Mitochondria play pivotal roles in axon degeneration, due to their importance in energy production, calcium homeostasis, production of reactive oxygen species and signaling [38, 39]. We transduced i3Neurons with a mitochondrial-targeted GFP and plated them in the center of wells to allow axons to project outwards and visualize individual axons. After axotomy, mitochondria underwent rapid shrinkage (Fig. 1d-g; Video S1) and increased in number (Fig. 1h). Using live imaging, we determined that the increase in mitochondrial number was due to fission events rather than mitochondrial transport and localization to the site of injury. Mitochondrial fission was most evident from close analysis of live imaging (Video S1 and Fig. 1d-e). This phenomenon was observed within 5 seconds after axon injury and was not affected by distance from the damage site from the cell body (Fig. S1a).

Mitochondrial fission is a dynamic process that is regulated by proteins localized in the inner and outer mitochondrial membranes [15]. A critical regulator of fission is dynamin related protein 1 (DRP1), a cytoplasmic GTPase. DRP1 is recruited to the outer mitochondrial membrane and helically oligomerizes around the mitochondrion to constrict it and bring about scission of the membrane. To test whether the fission we observed was dependent on DRP1, we knocked down DRP1 using CRISPR interference by expressing a previously validated guide RNA to DRP1 in i3Neurons expressing a catalytically dead Cas9 (dCAS9) [21, 37]. DRP1 knockdown prevented the increase in mitochondrial particle number (Fig. 1h). We confirmed this result by inhibiting DRP1 using P110, a peptide that blocks the interaction between DRP1 and FIS1 (mitochondrial fission protein 1), another protein that is essential for fission (Fig. 1i). In line with previous reports [40, 41], these events were dependent on calcium influx into the injured axon which, when buffered using BAPTA-AM, also prevented mitochondrial fission (Fig. S1a-d; Videos S2-5).

### Mitochondrial fission precedes proximal axon degeneration

To characterize the spatial and temporal dynamics of the axonal response to injury, we analyzed mitochondrial fission and axon degeneration in 100 µm segments of the axon relative to the injury site every 10 min following axotomy. This analysis revealed that mitochondrial fission began seconds after injury and progressed in a retrograde manner back towards the cell body. One hour after axotomy, mitochondria 500 µm away from the damage site had undergone fission (Fig. 1j-k, Video S6). This fission preceded a slower retrograde wave of axon degeneration back towards the cell body (Fig. 1l-m, Video S7).

Time lapse imaging of the cell bodies of axotomized neurons revealed that the retrograde wave of axon degeneration was followed by fragmentation of mitochondria in the soma and cell death (Fig. 1n-o; Videos S8-9). Axotomy led to death of 84% of cells analyzed. We have thus uncovered an exciting new human neuron model in which axon injury initiates progressive retrograde axon degeneration that ultimately results in neuronal cell death.

### DLK regulates axotomy-induced mitochondrial fission

To understand mechanisms governing neurodegeneration in this model, we started by examining a pathway involved in the cellular response to axon injury. The dual leucine zipper kinase (DLK; MAP3K12) is a key regulator of the axon damage response and neuronal cell death [29, 42]. Because mitochondria are critical regulators of axon degeneration, we were curious whether DLK protein might localize to axonal mitochondria. In i3Neurons, we observed association of endogenous DLK with 77% of axonal mitochondria (Fig S2a-c). We further corroborated this association by immuno-EM (Fig. S2d), and through observation of DLK co-trafficking with mitochondria (Fig. S2e).

To investigate how DLK responds to axon damage in human neurons, we performed laser axotomies and live imaging of GFP-tagged DLK and mitochondria. We observed an intriguing presence of DLK puncta at 45% of sites of mitochondrial fission (Fig. 2a-b, Video S10). To determine whether DLK influences mitochondrial fission, we generated DLK KO neurons using CRISPR/Cas9 (Fig. S2f). The knockout neurons were validated for absence of DLK protein by immunofluorescent staining and western blotting (Fig. S2g-h). In the undamaged condition, we observed no mitochondrial phenotypes in these cells including number, size, movement, and oxygen consumption rate (Fig. S2i-p). Strikingly, DLK KO neurons did not exhibit the axonal mitochondrial fission we had observed in response to axotomy in wild type neurons (Fig. 2c-d; Videos S11-12). Corroborating this result, pharmacological inhibition of DLK using GNE3511 prevented mitochondrial fission after injury (Fig. 2e, Videos S13-14).

**Figure 2.**
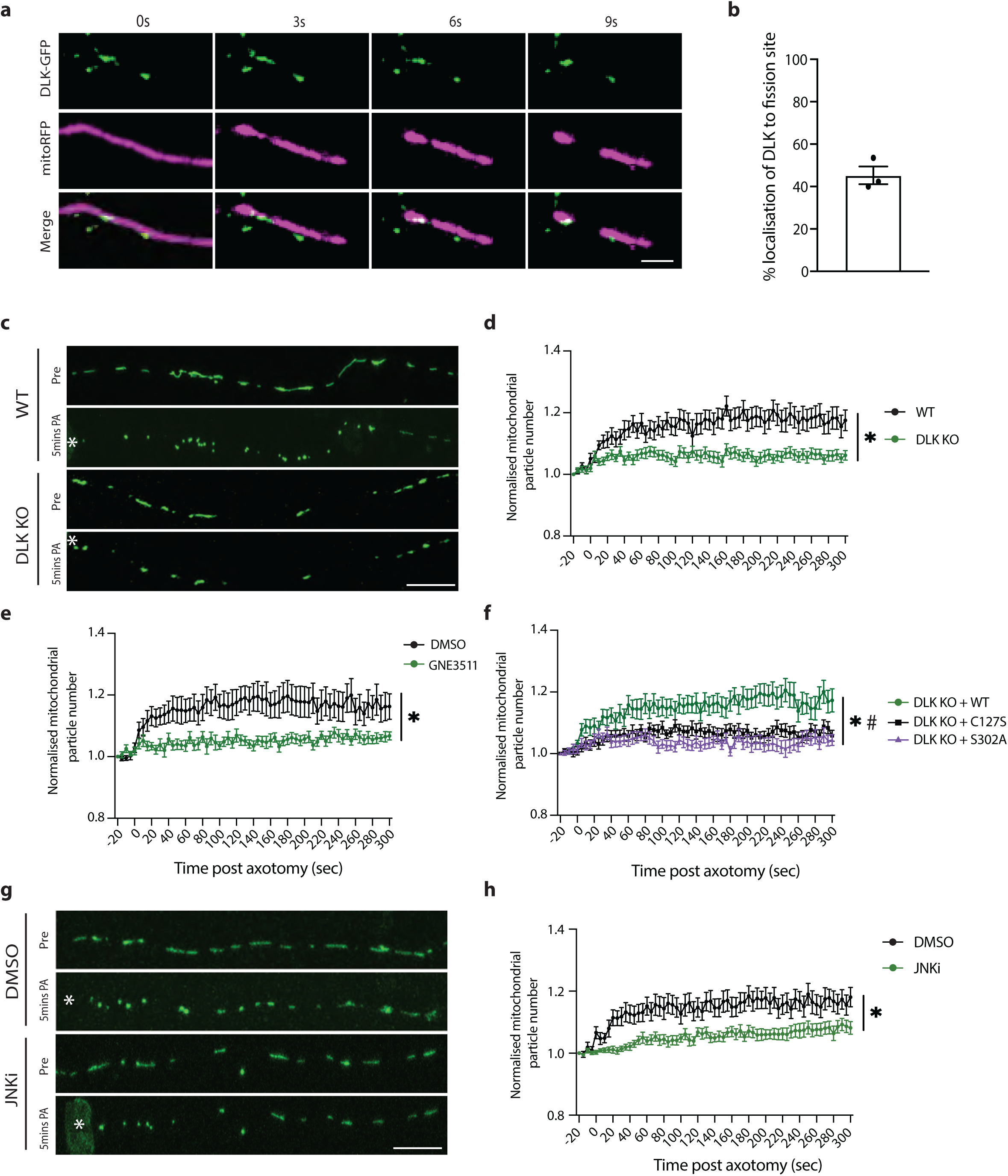
DLK regulates axotomy-induced mitochondrial fission. a) Representative images of DLK colocalization at the site of mitochondrial fission after axotomy. DLK-GFP (green) and mitochondria (mitoRFP, magenta). (Scale bar = 10 µm). b) Percentage mitochondrial fission events where DLK is localized at the site of fission. Results are represented as mean ± SEM. N=3 independent differentiations, N≥30 axotomized neurons. c) Representative images of WT and DLK KO neurons transduced with mitoGFP (green) pre (Pre) and 5 mins post axotomy (PA). * Marks the site of axotomy. (Scale bar = 25 µm). d) Normalized number of mitochondrial particles post axotomy in WT (black) and DLK KO (green) neurons. Results are represented as mean ± SEM. N=3 independent differentiations, N≥ 30 axotomized neurons. (Two-way ANOVA, Bonferroni correction (p≤0.05 *) e) Normalized number of mitochondrial particles post axotomy in DMSO (black) and GNE3511 (green)-treated neurons. Results are represented as mean ± SEM. N=3 independent differentiations, N≥ 25 axotomized neurons. (Two-way ANOVA, Bonferroni correction (p≤0.05 *) f) Normalized number of mitochondrial particles post axotomy in DLK KO neurons transduced with WT DLK-GFP (green), DLK-C127S-GFP (black) or DLK-S302A-GFP (purple). Results are represented as mean ± SEM. N=3 independent differentiations, N≥ 23 axotomized neurons. Two-way ANOVA, Bonferroni correction (*p≤0.05, WT DLK vs DLK C127S, # p≤0.05, WT DLK vs S302A). g) Representative images of neurons transduced with mitoGFP (green) treated with DMSO and JNKi pre (Pre) and 5 mins post axotomy (PA). * Marks the site of axotomy. (Scale bar = 25 µm). h) Normalized number of mitochondrial particles post axotomy in DMSO (black) and JNKi (green)-treated neurons. Results are represented as mean ± SEM. N=3 independent differentiations, N≥ 30 axotomized neurons. (Two-way ANOVA, Bonferroni correction (p≤0.05*)

To better understand the mechanism by which DLK regulates mitochondrial fission, DLK KO neurons were transduced with wild type DLK or 2 mutant versions of DLK. DLK C127S cannot be palmitoylated, hindering its ability to properly signal [43] . Kinase dead DLK (S302A) cannot be trans-phosphorylated and is catalytically inactive [27, 44]. Only the expression of wild type DLK rescued the fission phenotype in DLK KO neurons (Fig. 2f; Videos S15-17), indicating DLK kinase activity is critical for this process.

We next tested whether inhibiting the c-Jun N-terminal kinases (JNK), known downstream players of the DLK pathway, reduced mitochondrial fission after axotomy. Neurons exposed to JNK inhibitor (JNKi) showed a significant reduction in mitochondrial fission following axon injury (Fig. 2g-h). We thus uncover a role for DLK in initiating a wave of mitochondrial fission after injury.

### DLK kinase is necessary for DRP1 to induce mitochondrial fission

DRP1 phosphorylation state regulates its function, with phosphorylation at serine 616 promoting mitochondrial fission [45]. We speculated that DLK might be required for phosphorylation of DRP1 at this residue. In order to more easily study this signaling mechanism, we turned to a heterologous system, HEK293 cells, that do not express DLK. Overexpression of DLK in these cells can be used to study DLK signaling since its homodimerization is sufficient to drive pathway activity [44, 46].

Expression of wild type DLK, but not DLK C127S (palmitoyl-site mutant) or DLK S302A (kinase dead), resulted in activation of DLK signaling as reflected by the increased phosphorylation of its downstream target c-Jun (Fig. 3a-b). Interestingly, when we co-expressed DLK and DRP1, phosphorylation at S616 was increased in this model of DLK activation (Fig. 3a,c), but not with mutant DLK, showing that DLK activity is sufficient to cause DRP1 S616 phosphorylation. Levels of total DRP1 remained unchanged, as did levels of S637 pDRP1, a site whose phosphorylation is thought to impair DRP1 GTPase activity [47, 48] (Fig. 3a, d-e).

**Figure 3.**
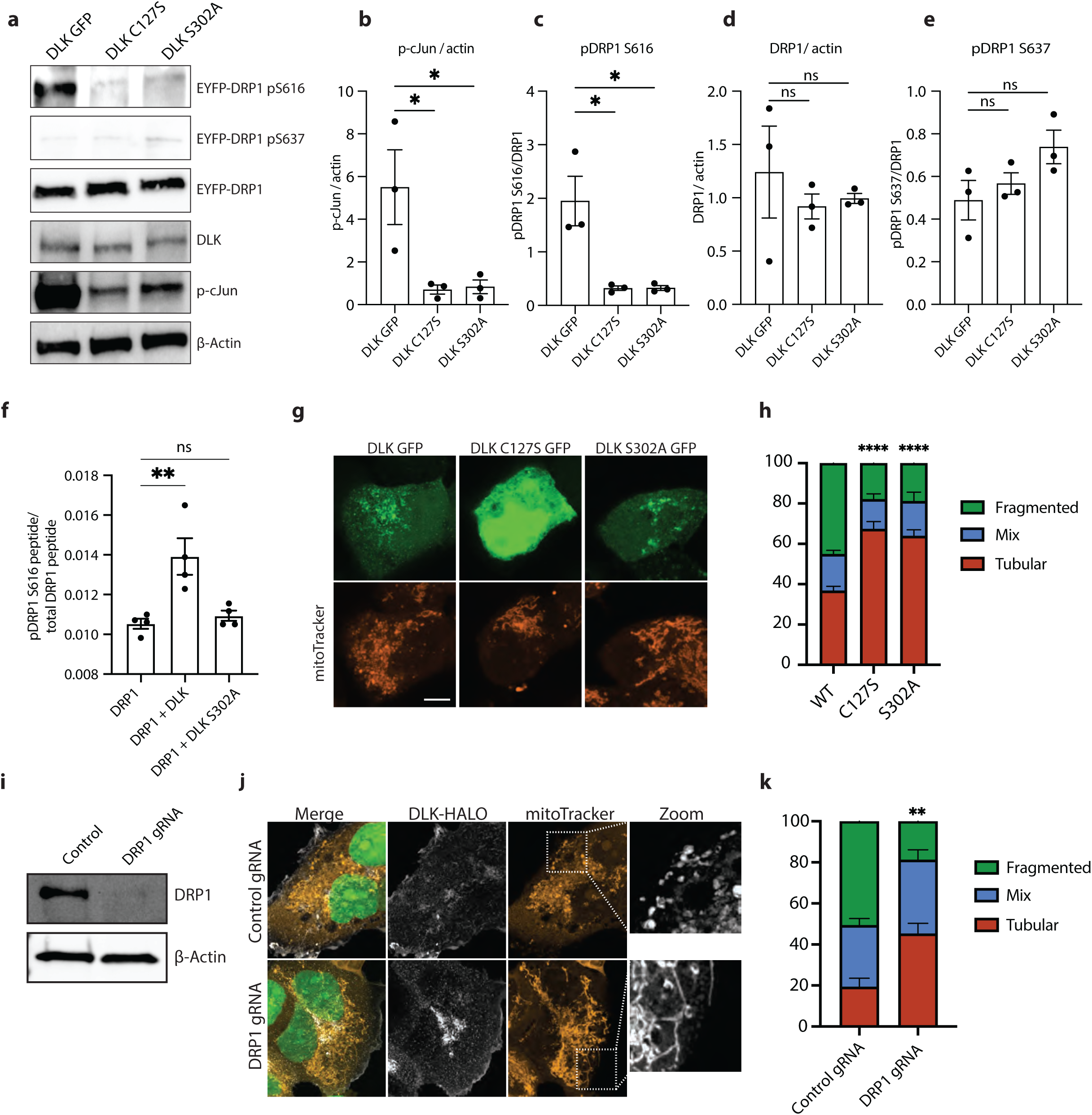
Activation of the DLK pathway by overexpression causes increased DRP1 phosphorylation. a) Representative western blots of HEK293A cells transfected for 24 hours with WT, C127S and S302A DLK-GFP. Immunoblot for pS616-DRP1, pS637-DRP1, total DRP1, DLK, pS63-cJun and loading control ß actin. b) Quantification of pS63-cJun/ ß actin levels after 24-hour expression of WT, C127S and S302A DLK-GFP in HEK293A cells. Results are represented as mean ± SEM. One-way ANOVA, Bonferroni correction (p≤0.05 *). c) Quantification of pS616-DRP1/total DRP1 levels after 24-hour expression of WT, C127S and S302A DLK-GFP in HEK293A cells. Results are represented as mean ± SEM. One-way ANOVA, Bonferroni correction (p≤0.05 *). d) Quantification of total DRP1/ ß actin levels after 24-hour expression of WT, C127S and S302A DLK-GFP in HEK293A cells. Results are represented as mean ± SEM. One-way ANOVA, Bonferroni correction (not significant, ns). e) Quantification of pS637 DRP1/ ß actin levels after 24-hour expression of WT, C127S and S302A DLK-GFP in HEK293A cells. Results are represented as mean ± SEM. One-way ANOVA, Bonferroni correction (not significant, ns). f) Phospho-mass spectrometry. pS616-DRP1 peptide / total DRP1 peptide levels from purified GST-DRP1 in HEK 293A cells expressing GST-DRP1, GST-DRP1 and DLK-GFP or GST-DRP1 and DLK-S302A-GFP. Results are represented as mean ± SEM. One-way ANOVA, Bonferroni correction (p≤0.01 **). g) Representative images of HEK-293A cells expressing WT, C127S and S302A DLK-GFP (green) and mitoTracker (orange) for 24 hours. (Scale bar = 10 µm). h) Quantification of mitochondrial morphology in HEK293A cells transfected for 24 hours with WT, C127S and S302A DLK-GFP. N=3 individual transfections. Results are represented as mean ± SEM. Two-way ANOVA, Bonferroni correction (WT vs C127S p≤0.0001 ****, WT vs S302A p≤0.0001 ****, C127S vs S302A not significant, ns). i) Representative western blots HEK-293A cells expressing dCAS9 transduced with control and DRP1 gRNAs. Immunoblot for total DRP1 and loading control ß actin. j) Representative images of Control and DRP1 KD HEK-293A cells expressing DLK-HALO for 24 hours. NLS-GFP (green), DLK-HALO (white), mitoTracker (orange). (Scale bar = 10 µm). k) Quantification of Control and DRP1 KD HEK-293A cells mitochondrial morphology. Cells transfected for 24 hours with DLK-HALO. N=3 individual transfections. Results are represented as mean ± SEM. (Two-way ANOVA, Bonferroni correction (p≤0.01 **)

To further validate that DLK signaling causes DRP1 phosphorylation, we purified GST-DRP1 from HEK293 cells expressing wild type DLK or DLK S302A, and performed targeted mass spectrometry to quantify DRP1 phosphorylation. The relative abundance of phospho-DRP1 S616 peptides was significantly increased only in the HEK293 cells expressing wild type but not kinase dead DLK (Fig. 3f). These results demonstrate that DLK kinase activity can promote DRP1 phosphorylation at S616.

To further examine this phosphorylation event, we performed an *in vitro* kinase assay using proteins separately purified from HEK293 cells. Incubation of DRP1 with wild type DLK increased pDRP1 S616, whereas incubation with kinase dead DLK did not (Fig. S3a-c). We wondered whether this DLK kinase activity on DRP1 was occurring via the canonical DLK kinase cascade, since purification of DLK from HEK cells could result in isolation of the entire DLK signaling complex including downstream kinases MAP2K and JNK. We detected JNK to be associated with the DLK we pulled down, suggesting this was via canonical MAP3K signaling (Fig S3d). We also found that DLK inhibitor and JNK inhibitor (but not p38 inhibitor) could block the phosphorylation of DRP observed in the heterologous expression of DLK in HEK cells (Fig S3d-f). Together, these data demonstrate that DRP1 is a novel substrate of DLK signaling that occurs via the canonical DLK kinase cascade. To our knowledge, DRP1 is only the second substrate of DLK/JNK signaling to be defined after cJun.

We analyzed mitochondrial morphology in the HEK293 cells to examine whether DRP1 phosphorylation caused by heterologous expression of DLK in HEK cells led to fission. Similar to what we had observed in injured axons, DLK signaling resulted in an increase in mitochondrial fragmentation (Fig. 3g-h). DLK-induced mitochondrial fragmentation was prevented by knocking down DRP1 (Fig. 3i-k). These data identify DLK as a novel kinase upstream of DRP1 whose role is to drive mitochondrial fission via phosphorylation of DRP1 at serine 616.

### Axotomy causes DLK-dependent phosphorylation of DRP1

To validate these results in the i3Neurons, DLK KO or WT neurons were center-plated such that axon fractions could be harvested to examine axonal DRP1 phosphorylation after injury (Fig. 4a). Two hours after axotomy, phospho-DRP1 S616 (pDRP1 S616) was increased in axons but not cell bodies of wild type neurons. However, as predicted, the increase in p-DRP1 did not occur in DLK KO axons (Fig. 4b-e). The increase in pDRP1 S616 in wild type but not DLK KO axons was further validated by immunostaining (Fig. 4f-g).

**Figure 4.**
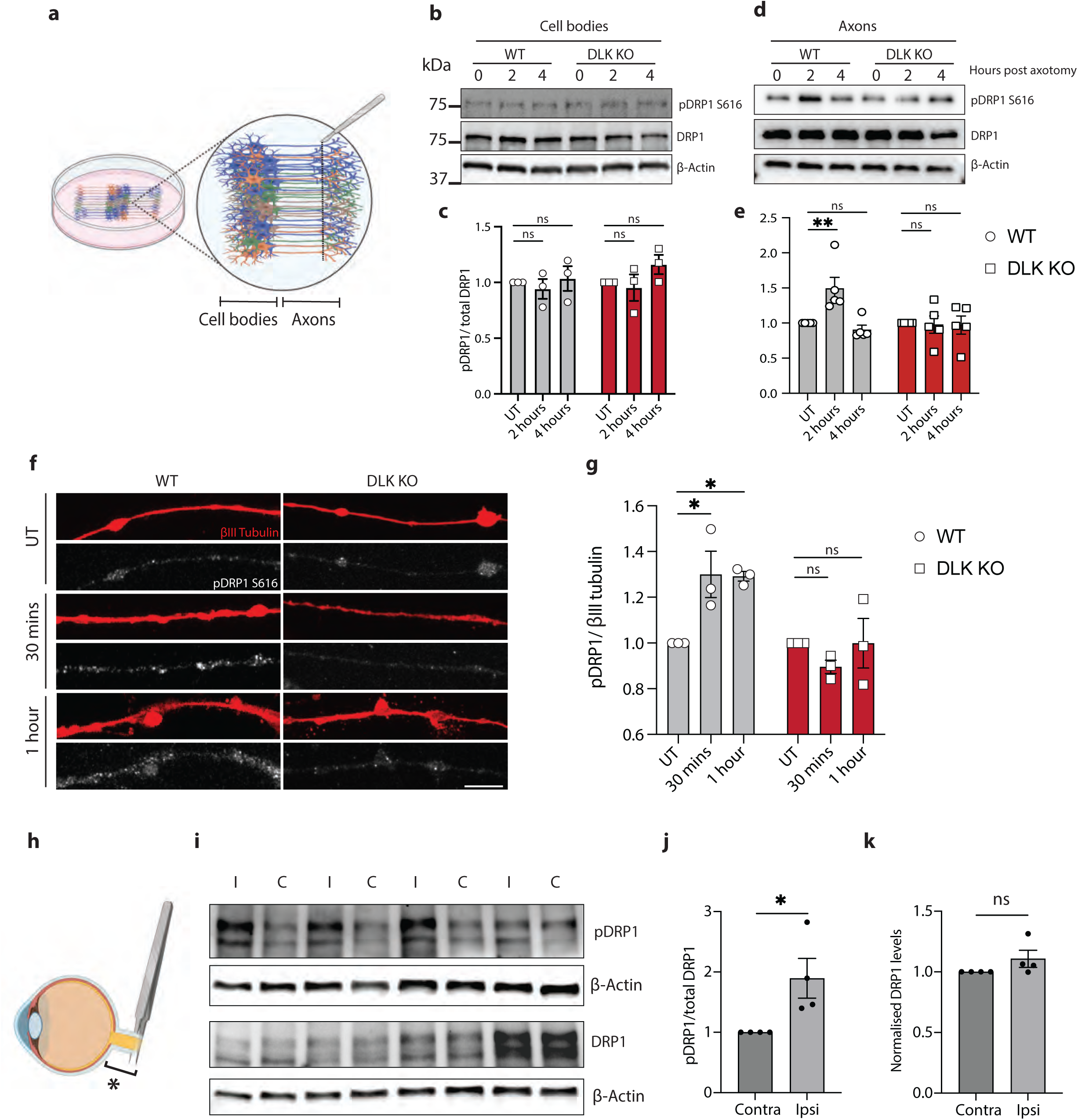
Axotomy causes DLK-dependent phosphorylation of DRP1. a) Schematic representation of i3Neuron center plating and axotomy for protein harvesting after injury. b) Representative western blots of WT and DLK KO neuron cell bodies untreated (UT), 2 and 4 hours post axotomy. Immunoblot for p-S616 DRP1, total DRP1 and loading control ß actin. c) Quantification of pDRP1 (S616)/ total DRP1 levels in WT and DLK KO neuron cell bodies untreated (UT), 2 and 4 hours post axotomy. Results normalized to UT. Results are represented as mean ± SEM. N=3 independent differentiations. One-way ANOVA. No significant changes observed. d) Representative western blots of WT and DLK KO neuron axons untreated (UT), 2 and 4 hours post axotomy. Immunoblot for p-S616 DRP1, total DRP1 and loading control ß actin. e) Quantification of pDRP1 (S616)/ total DRP1 levels in WT and DLK KO neuron axons 0, 2 and 4 hours post axotomy. Results normalized to UT. Results are represented as mean ± SEM. N=3 independent differentiations. Two-way ANOVA. Bonferroni correction (p≤0.05*). f) Representative images of WT and DLK KO axons stained for ßIII tubulin (red) and pDRP1 S616 (white) untreated (UT), 30 mins and 1 hour post axotomy. (Scale bar = 25 µm). g) Quantification of pDRP1 S616 fluorescence in WT and DLK KO neuron axons after axotomy. Results normalized to untreated axons. Results are represented as mean ± SEM. N=3 independent differentiations. Two-way ANOVA, Bonferroni correction (p≤0.05 *) h) Schematic representation of optic nerve crush injury model. Bracket and * indicate the proximal portion of the injured nerve harvested for western blotting. i) Representative western blots of proximal portion of ipsilateral (I) injured and contralateral (C) optic nerves 24 hours after optic nerve crush. Immunoblot for p-S616 DRP1, total DRP1 and loading control ß actin. j) Quantification of pDRP1 (S616)/ total DRP1 levels after optic nerve crush. Results normalized to contralateral side (C). Results are represented as mean ± SEM. Unpaired t-test (p≤0.05 *). k) Quantification of total DRP1/ ß actin levels after optic nerve crush. Results normalized to contralateral side (C). Results are represented as mean ± SEM. Unpaired t-test not significant, ns.

To examine whether DRP1 phosphorylation at S616 also occurs *in vivo*, we performed optic nerve crush in mice. Increased pDRP1 S616 relative to total DRP1 was detected in optic nerve proximal to the crush site 24 hours after injury (Fig. 4h-j). Levels of total DRP1 in injured nerves remained unchanged (Fig. 4i,k).

### Inhibiting DRP1 delays axon degeneration and neuron death

Since axotomy leads to progressive degeneration of the axon proximal to the injury and eventually to cell death, we asked to what extent DLK and DRP1 control the degeneration of each compartment. DLK has been shown to contribute to axon degeneration and to neuron death in many contexts [26–30, 42]. We hypothesized that the DLK pathway would promote axon degeneration in axotomized human neurons. Whereas wild type axons had degenerated as early as 4 hours after injury, DLK KO axons were still present after 24 hours (Fig. 5a-c). Axons of neurons treated with JNK inhibitor were similarly protected (Fig S4c-d). Interestingly, in neurons with CRISPRi knockdown of DRP1, axons were also protected from degeneration (Fig. 5d-f).

**Figure 5.**
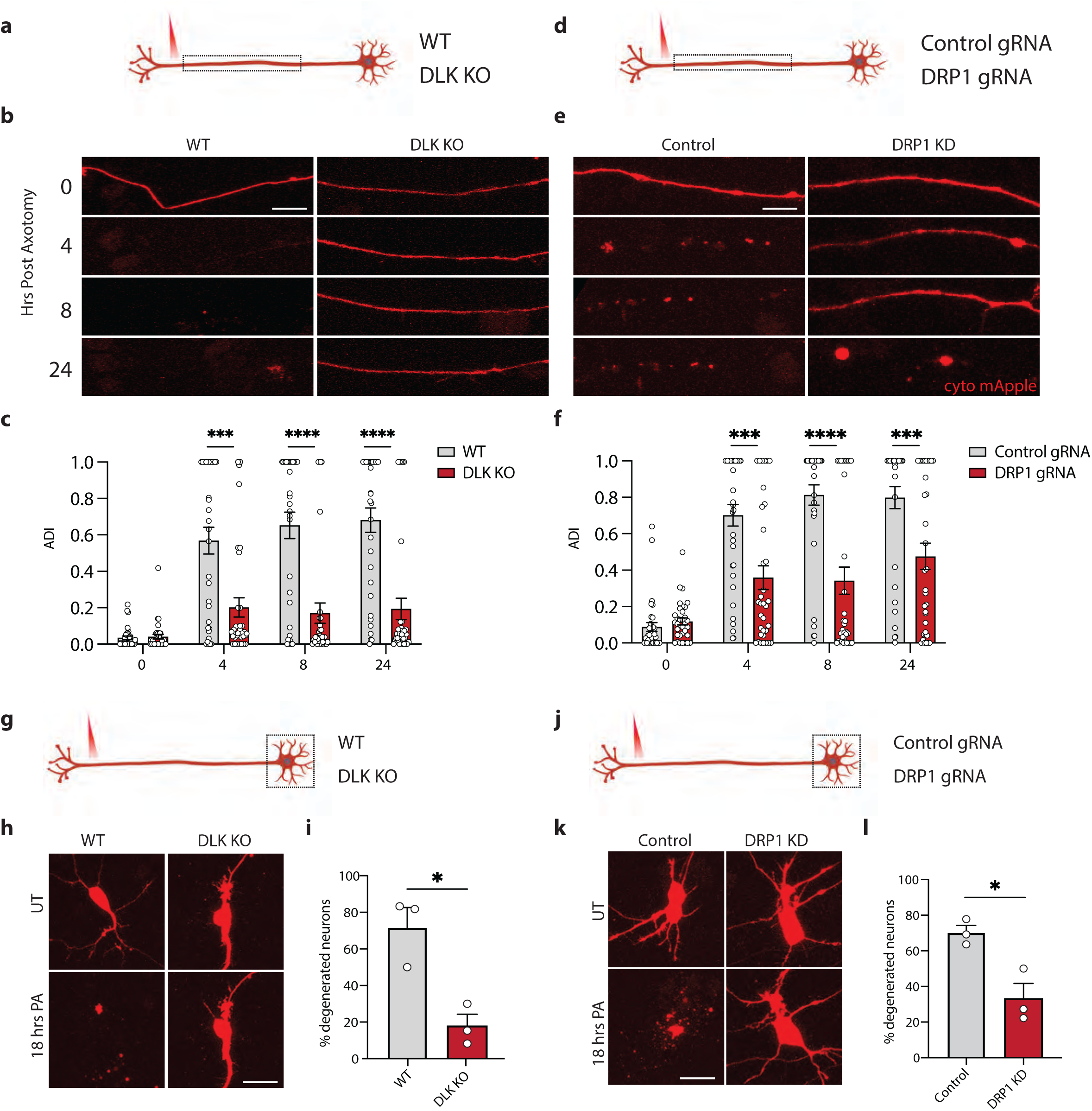
Blocking DRP1 or DLK delays axon degeneration. a) Schematic representation of i3Neuron laser axotomy. Dashed box indicates the field of view b) Representative images of WT and DLK KO neuron axons transduced with cyto mApple (red) proximal to the site of injury 0,4, 8 and 24 hours post axotomy (Scale bar = 25 µm). c) Quantification of axon degeneration index (ADI) in WT and DLK KO neurons 0, 4, 8 and 24 hours post axotomy. Results are represented as mean ± SEM. N=3 independent differentiations, N≥ 33 axons. Two-way ANOVA, Bonferroni correction (p≤0.005 ***, p≤0.001****). d) Schematic representation of i3Neuron laser axotomy. Dashed box indicates field of view. e) Representative images of Control and DRP1 KD neuron axons transduced with cyto mApple (red) proximal to the site of injury 0, 4, 8 and 24 hours post axotomy (Scale bar = 25 µm). f) Quantification of axon degeneration index (ADI) in WT and DLK KO neurons 0, 4, 8 and 24 hours post axotomy. Results are represented as mean ± SEM. N=3 independent differentiations, N≥ 33 axons. Two-way ANOVA, Bonferroni correction (p≤0.005 ***, p≤0.001****). g) Schematic representation of i3Neuron laser axotomy. Dashed box indicates field of view. h) Representative images of WT and DLK KO neuron cell bodies transduced with cyto mApple (red) pre and 18 hours post axotomy (PA). (Scale bar = 40 µm). i) Percentage degenerated WT and DLK KO 18 post axotomy. Results are represented as mean ± SEM. N=3 independent differentiations. Unpaired t-test (p≤0.05 *). j) Schematic representation of i3Neuron laser axotomy. Dashed box indicates field of view. k) Representative images of Control and DRP1 KD neuron cell bodies transduced with cyto mApple (red) pre and 18 hours post axotomy (PA). (Scale bar = 40 µm). l) Percentage degenerated Control and DRP1 KD 18 post axotomy. Results are represented as mean ± SEM. N=3 independent differentiations. Unpaired t-test (p≤0.05 *).

To determine whether loss of DLK or DRP1 also protected neurons from death, we followed the fate of cell bodies of axotomized neurons. In control neurons we observed cell death in about 70% of cells after axotomy whereas only 18% of DLK KO cells and 33% of DRP1 knockdown cells died (Fig 5g-l). These results demonstrate that both DRP1 and DLK are key regulators of neuronal cell death after axotomy. Interestingly, inhibiting transcription using actinomycin D (ActD) protected neurons from cell death but did not prevent the retrograde axon degeneration (Fig S4a-b), suggesting that DLK drives neuronal degeneration via distinct mechanisms depending on the compartment.

### Loss of DRP1 prevents neuron death after axotomy *in vivo*

We then asked how universal this pathway is, or whether it was unique to the i3Neuron model. It has been established that silencing DLK blocks retinal ganglion cell (RGC) death after optic nerve crush [27, 28, 33, 36]. However, DRP1 has not previously been implicated downstream of this DLK signaling. To knock down DRP1, we used an AAV delivery strategy to express a small Cas9 (sa-Cas9) together with a guide RNA to DRP1 [49] (Fig 6a). AAVs were injected into the intravitreal space to transduce RGCs. Three weeks after the injection, DRP1 protein levels were significantly reduced in the optic nerve of mice that had received gRNA DRP1 but not control scrambled gRNA, validating this approach (Fig 6b-c).

**Figure 6.**
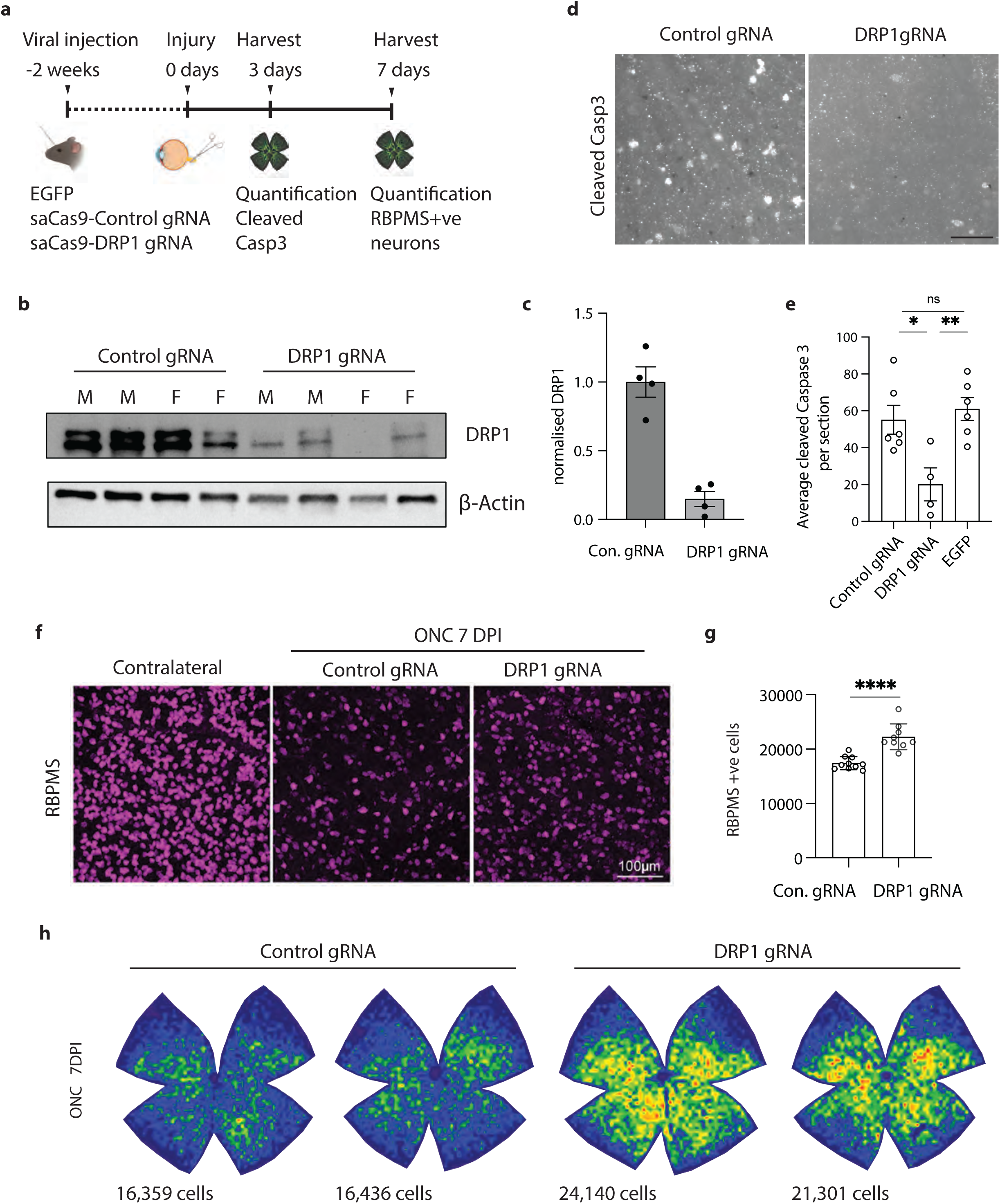
DRP1 mediates cell death after ONC. a) Schematic representation of RGC survival following optic nerve crush (ONC). b) Western blots of Control and DRP1 gRNA transduced optic nerved. Immunoblot for DRP1 and loading control ß actin. c) Quantification of DRP1 knockdown by Western blot. Results are represented as mean ± SEM. d) Representative images of cleaved caspase 3 positive cells in saCas9-Control and saCas9-DRP1 gRNA transduced RGCs 3 days after ONC. (Scale bar = 25 µm). e) Average cleaved caspase 3 positive cells per section in saCas9-Control, saCas9-DRP1 gRNA and EGFP transduced RGCs after nerve crush. Results are represented as mean ± SEM. N≥4 animals per condition. One-way ANOVA, Bonferroni correction. (Control gRNA vs EGFP, not significant, ns. Control gRNA vs DRP1 gRNA p≤0.05 *. DRP1 gRNA vs EGFP p≤0.01 **). f) Representative images of retinas 7 days post ONC and immunostained for RGC marker RBPMS (magenta). Images acquired at similar areas and the same distance from the optic nerve head (ONH). Scale bar = 100 µm. g) Quantification of RBPMS-positive RGCs in the retinas of control and DRP1 gRNA mice at 7DPI. (N=8 mice per condition. Unpaired t-test P < 0.0001 ****) h) Representative isodensity maps display the topographical survival of RBPMS+RGCs at 7 days post-injury. DRP1 gRNA treatment delays ONC-induced RGC degeneration across the retina. Color scale for isodensity maps ranges from 0 (purple) to 3,600 (red) RGCs/mm².

Optic nerve crush was performed two weeks after AAV injection, and retinas were harvested 3 days post injury, a time point at which injured RGCs express cleaved caspase 3 (Fig. 6a). The number of cleaved caspase 3-positive RGCs with saCas9 and a control gRNA was similar to a control virus expressing just EGFP (55 ± 7.83 and 61 ± 6.24, respectively; average caspase-positive cell count per ROI +/- SEM). However, knocking down DRP1 significantly reduced the caspase positive cell count to 20 ± 8.93 (Fig. 6d-e), showing that apoptosis of RGCs relies on DRP1.

We performed a parallel experiment with the same viral strategy for DRP1 knockdown to assess RGC survival 7 days after the injury. Using the RGC marker RBPMS, we found that DRP1 knockdown protected RGCs from axotomy-induced death (Fig 6f-h), increasing the total RGC number per retina from 17,423 ± 379 to 22,273 ± 789 (average +/- SEM). Together, these experiments demonstrate that DRP1 contributes to neuronal cell death after axon injury *in vivo*.

### BAX mediates axon degeneration and cell death after axotomy

To better understand the type of axon degeneration and cell death observed in our study, we asked whether they occurred via canonical pathways of apoptosis, and in particular whether they involved BAX. In models of apoptosis, DRP1 contributes to BAX recruitment to mitochondria [50], and DRP1 and BAX interact to drive mitochondrial depolarization and fragmentation [19, 51]. Jenner et al. showed a close interaction between DRP1 and BAX in HEK cells undergoing apoptosis using a fluorescence complementation biosensor (ddRFP [52]). This system produces enhanced red fluorescence upon dimerization of A_1_ and B_1_ proteins. Using A_1_-DRP1 and B_1_-BAX fusion constructs, we tested whether DRP1 and BAX closely interact in the axon after injury. One hour after axotomy, we observed a significant increase in red fluorescence. The interaction between DRP1 and BAX occurred at mitochondria as shown by co-localization with mito-GFP (Fig. 7a). However, DLK inhibitor GNE3511 reduced their interaction (Fig. 7a-b). To test whether the interaction between DRP1 and BAX was dependent on DRP1 phosphorylation at S616, we mutated the B_1_-DRP construct to carry a S616A point mutation, preventing its phosphorylation at this site. A_1_-BAX and B_1_-DRP1 S616A did not show the increase in red fluorescence, demonstrating that the BAX-DRP1 interaction is more likely to occur if DRP1 is phosphorylated at S616 (Fig. 7c), which requires DLK (Fig. 3).

**Figure 7.**
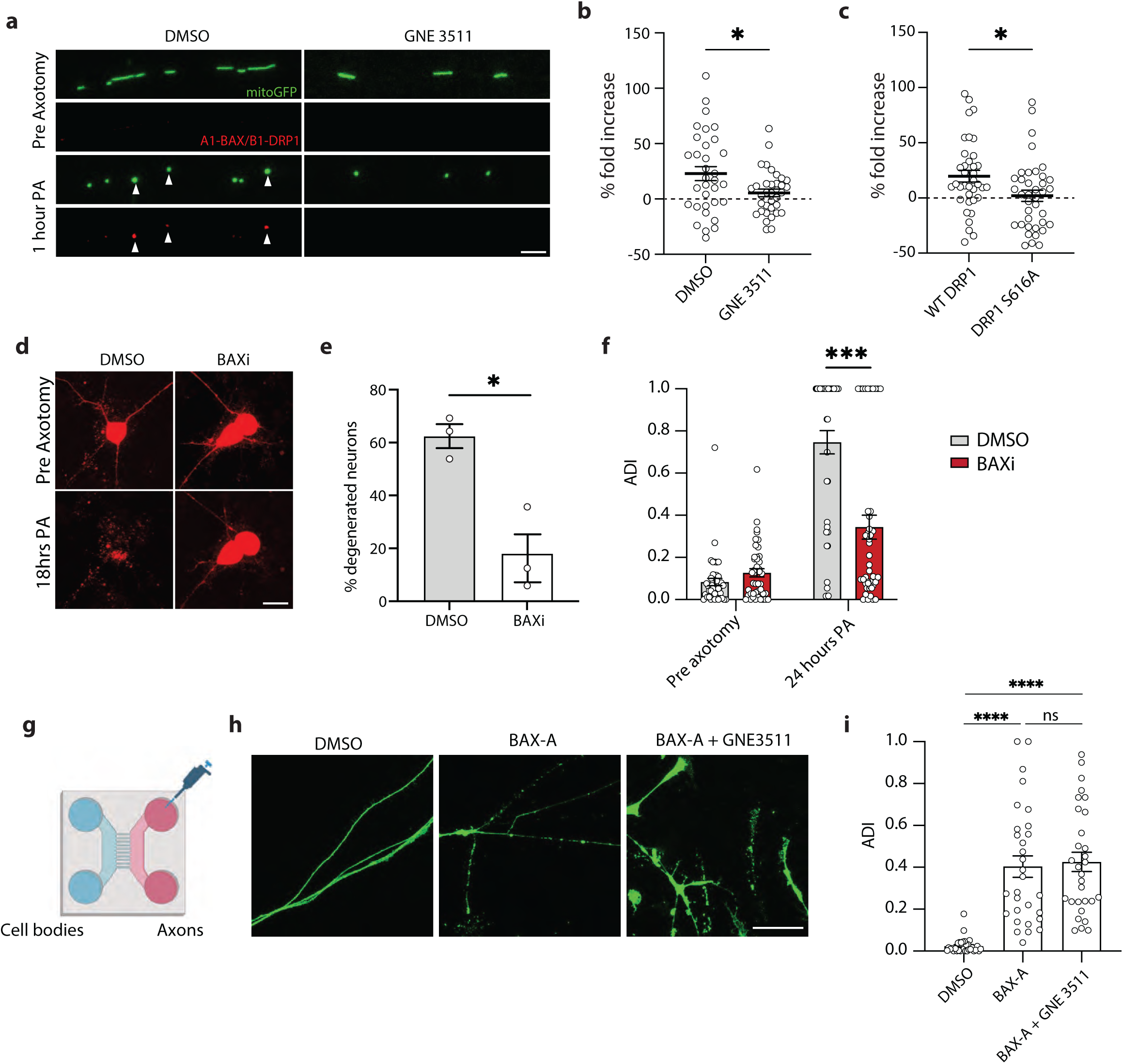
BAX regulates cell death after axotomy. a) Representative images of WT neurons treated with DMSO or GNE3511 pre- and 1-hour post- axotomy (PA) expressing mitoGFP (green) and A_1_-BAX/B_1_-DRP1 complexes (red). (Scale bar = 5 µm). b) Quantification of A_1_-BAX/B_1_-DRP1 fluorescence fold change in the mitochondria of DMSO or GNE3511 treated neurons 1-hour post-axotomy. Results are represented as mean ± SEM. N=3 independent differentiations, N≥ 34 axotomized neurons. Two-way ANOVA, Bonferroni correction. (p≤0.05 *). c) Quantification of A_1_-BAX/B_1_-WT DRP1 and A_1_-BAX/B_1_-DRP1 S616A fluorescence fold change in the mitochondria 1-hour post-axotomy. Results are represented as mean ± SEM. N=3 independent differentiations, N≥ 34 axotomized neurons. Two-way ANOVA, Bonferroni correction. (p≤0.05 *). d) Representative images of DMSO and BAXi treated neuron cell bodies transduced with cyto mApple (red) pre and 18 hours post axotomy (PA). (Scale bar = 40 µm). e) Percent degenerated neurons 18 hours post axotomy treated with either DMSO or BAXi. Results are represented as mean ± SEM. N=3 independent differentiations. Unpaired t-test (p≤0.05 *). f) Quantification of axon degeneration index (ADI) of DMSO and BAX-A-treated neurons 24 hours after axotomy. Results are represented as mean ± SEM. N=3 independent differentiations. One-way ANOVA, Bonferroni correction (p≤0.001 ***, not significant, ns). g) Illustration of microfluidic devices used to separate axons from somas allowing for the local treatment of i3Neuron axons. h) Representative images of WT axons separated using microfluidic chambers treated with DMSO, BAX-A or BAX-A + GNE3511 for 24 hours. (Scale bar = 25 µm). i) Quantification of axon degeneration index (ADI) of isolated axons treated with DMSO BAX-A and BAX-A + GNE3511 for 24 hours. Results are represented as mean ± SEM. N=3 independent differentiations. One-way ANOVA, Bonferroni correction (p≤0.001 ****, not significant, ns).

We next tested whether inhibiting BAX would prevent cell death and axon degeneration in i3Neurons using BAI, a highly selective small molecule inhibitor of BAX [53]. BAX inhibition significantly preserved cell bodies of axotomized neurons (Fig. 7d-e). BAX inhibition also significantly reduced axon degeneration measured 24 hours after injury (Fig. 7f), indicating that BAX is a key mediator of axon degeneration and cell death after axotomy.

Finally, we asked whether local activation of BAX in the axon was sufficient to cause axons to degenerate. Other studies have shown stimulation of BAX with a Bcl-x antagonist causes axon degeneration [3, 54]. We grew i3Neurons in microfluidic chambers to physically isolate axons from cell bodies and dendrites. BAX activator (BAX-A), which binds to the N-terminus of BAX to promote its oligomerization [55], was added to the axon compartment, and we measured axon integrity 24 hours later. We also performed this experiment in the presence of DLK inhibitor GNE3511 (Fig. 7h). Local activation of BAX caused axons to degenerate, independently of DLK inhibition (Fig. 7h-i), suggesting BAX signaling is downstream of DLK.

Together, our data reveal a local DLK signaling cascade within the axon that drives retrograde axon degeneration after damage and does not require transcription. Starting at the site of damage, the DLK/JNK complex locally phosphorylates DRP1, recruiting BAX to mitochondria and producing an apoptotic cascade that spreads from the injury site back to the cell body where it eventually results in cell death.

## Discussion

In this work, we establish a novel and reproducible model of progressive axon degeneration and neuronal death in human neurons and unravel a cascade of events governing neurodegeneration after axon injury. We show that DLK kinase is required for DRP1 to drive mitochondrial fission and BAX-dependent apoptosis that originates in the axon and leads to degeneration of the axon followed by the cell body. We demonstrate the relevance of this pathway *in vivo*, extending from cultured human neurons to retinal ganglion cells of mice with optic nerve crush injuries. We hypothesize this pathway is broadly implicated in neuronal degeneration caused by axon damage, such as spinal cord injury and traumatic brain injury.

While BAX and DLK have separately been implicated in neuronal apoptosis of retinal ganglion cells after optic nerve crush [33, 36, 56, 57], an interaction between the two has not previously been reported, nor with DRP1. One study reported that DRP1-dependent mitochondrial fission occurs in axons after damage [58]. Here, we show how DLK, DRP1, and BAX cooperate to cause neuronal apoptosis after axon damage, and in particular, how the same apoptotic mechanism can follow axon loss [59] .

Until now, it was also unclear whether pathways distinct from SARM1 can mediate axon degeneration. Many clinical descriptions of axon degeneration refer to “dying back” as a pattern in which distal axons are lost before cell bodies [60, 61]. It has been suggested that dying back axon degeneration overlaps with Wallerian degeneration [62], but this remains unclear [63]. Although axon degeneration is rescued by SARM1 deletion in some models (e.g. chemotherapy-induced peripheral neuropathy [64]; TDP43-linked ALS [65]), it is not in others (SOD1 mouse model of ALS [7]; and optic nerve crush [8]). We have shown that a type of non-Wallerian axon degeneration that is more akin to apoptosis can occur after axon damage, and that it spreads retrogradely, similar to a dying back. It is possible that additional axon degeneration pathways also exist.

Therapeutically targeting this pathway offers multiple options. There is precedence for testing some of these strategies, such as blocking BAX [56, 57], or targeting mitochondrial dynamics [66], including promoting mitochondrial fusion [34] and blocking mitochondrial fission [67–70]. Blocking DLK signaling using small molecules has been considered [29, 32, 71], although a Phase I clinical trial in ALS patients was recently halted due to safety concerns [72]. However, specifically targeting the interaction between the DLK signaling complex and DRP1 has never been attempted.

In addition to protecting axons as we have shown, targeting DRP1 has the potential to protect dendrites [73]. However, it also has possible toxicity issues as mitochondrial fission is critical for regeneration and motor neuron survival after sciatic nerve injury [74]. A more complete understanding of DRP1 regulation in homeostatic and injury conditions will be helpful. For example, future studies may identify other targetable DRP1 phosphorylation sites in addition to serine 616 as well as additional kinases. To date, kinases of DRP1 that have been identified include CDK5 [75, 76], TBK1 [77], ERK1/2 [78, 79], CAMKIa [80] and CAMK2 [81]; here, we highlight DLK/JNK as a novel regulatory kinase upstream of DRP1 [82].

As a cytoplasmic facing kinase complex, DLK/JNK has the potential to regulate many substrates of which we now identify DRP1. Additional substrates of the pathway will be of great interest to uncover, as DLK is reported to associate with the plasma membrane, Rab3, Rab6, Rab7, Rab11, Golgi-derived, and LAMP1-positive vesicles [43, 83]. Localization of DLK to mitochondria has also recently been described in cultured mouse embryonic DRG neurons [83], corroborating our observation of DLK-mitochondrial association.

Our study highlights a novel signaling pathway downstream of DLK. It has generally been thought that DLK-dependent transcriptional events govern neuronal degeneration [36], but here we find that transcription is dispensable for retrograde axon degeneration driven by DLK via DRP1. It will be of interest to examine how the retrograde axon degeneration mechanism regulated by DLK-DRP1 is coordinated with transcriptional regulation of cell death factors by DLK, for example BID and PUMA [3].

DRP1-dependent apoptosis has been described in many cell types [17] however, its regulation by DLK was not previously recognized. That axonal DLK can locally trigger this cellular apoptotic pathway from a site of axonal injury that then spreads retrogradely to the cell body is distinct from the previously recognized transcriptional regulation of apoptosis by DLK. Our results call to mind new questions, such as how is DLK localized to the site of mitochondrial fission so it can regulate DRP1 phosphorylation? Does it associate with factors like MFF, which is an essential factor in recruiting DRP1 to mitochondria [84]? And what is the mechanism of activation of DLK that would link it to mitochondrial fission? In the future, it will be of great interest to explore under which conditions this pathway is activated, as well as the specificity of this pathway to different neuronal subtypes.

## Methods

### I3Neuron differentiation

I3Neurons were differentiated as previously described. Briefly, i^3^ iPSCs were dissociated using Accutase (Life Technologies, A1110501). Cells were plated in Matrigel-coated (1:100 Corning) plates in Neuronal induction media on day 0 (Knockout Dulbecco’s modified Eagle’s medium (DMEM)/F12 medium; Life Technologies Corporation, 12660012), 1X N2 supplement (Life Technologies, 17502048), 1× GlutaMAX (Thermofisher Scientific, 35050061), 1× MEM nonessential amino acids (NEAA) (Thermofisher Scientific, 11140050), 10 μM ROCK inhibitor (Y-27632; Tocris, 1254), and 2 μg/ml doxycycline (Clontech, 631311). Neuronal induction media was changed once a day for 2 more days. On day 3 of induction, cells were dissociated using Accutase and plated in dishes coated with poly-L-ornithine (PLO; 0.1 mg/ml; Sigma, P3655-10MG). Cells were plated in neuronal maturation media (BrainPhys medium (STEMCELL Technologies, 05790), 1× B27 Plus Supplement (ThermoFisher Scientific, A3582801), 10 ng/ml BDNF (PeproTech, 450-02), 10 ng/ml NT-3 (PeproTech, 450-03), 1 mg/ml mouse laminin (Invitrogen, 23017015), and 2 μg/ml doxycycline). Half of the neuronal maturation media was removed and replenished with fresh media every 2-3 days.

### Lentivirus generation

Lentivirus were generated as previously described [37], 7 million Lenti-X HEK-293T cells (Takara Bio, 632180) were seeded in 9 mL DMEM. The next day, a transfection mix was prepared containing 1 µg of Lenti plasmid, 3 µg of third generation packaging mix (1:1:1 mix of three plasmids), 12 µL Lipofectamine 3000 Reagent (ThermoFisher, L3000008), and 250 µL Opti-MEM I Reduced Serum Medium (GIBCO, 31985070). The mix was vortexed, spun down briefly, incubated at room temperature for 40 minutes, then added dropwise to the HEK cells and gently swirled to mix. The next day, the media was replaced with 18 ml fresh 10% FBS DMEM supplemented with 1:500 ViralBoost (Alstem, VB100). Two days later, the media was collected into a 50 ml Falcon tube, supplemented with 6 ml Lenti-X Concentrator (Takara Bio; 631231), mixed thoroughly, and stored at 4°C for 48 hours. The supernatant was then spun down at 4°C for 45 minutes at 1,500 x g. The supernatant was aspirated, and the pellet was resuspended in 500 μL of PBS.

### Plasmids and cloning

All DLK (Ensembl CCDS: 55831) constructs and mutants were generated by Epoch Live Science (Texas, USA) and cloned into a lentivirus-expressing backbone under a doxycycline-dependent promoter. DLK was C-terminally tagged by inserting a glycine rich linker (3x GGGGS) followed by EGFP, just before the stop codon. EYFP-DRP1 was obtained from Addgene (45160). For GST-purification, DLK and DRP1 were N-terminally tagged into a GST-containing backbone using In-fusion cloning (Takara Bio. 638945). Cyto-mAPPLE plasmid was a gift from Michael E. Ward (NINDS). A_1_-BAX and B_1_-DRP1 were gifted by Dr. Ana J. Garcia-Saez (Max Plank Institute of Biophysics). Mito-EGFP and Mito-RFP plasmids were gifts from Dr. Zu-Hang Sheng (NINDS).

### DRP1 knockdown using dead Cas9

For DRP1 knockdown, we used CRISPRi-i^3^ iPSCs containing a CAG promoter-driven dCas9-BFP-KRAB cassette inserted into the CLYBL safe harbor locus (Addgene #127968). iPSCs were transduced with control gRNA or a previously validated gRNA targeting DRP1^84^ and differentiated as described above.

### Laser Axotomies

Laser axotomies were performed on 10-14 day old neurons using a Zeiss LSM 880 Upright 2-Photon microscope equipped with an ablate laser. Temperature and CO_2_ levels were maintained using a Pecon microscope incubator. Ablation laser was set to 800 nm, 100% power and 100 iterations. Neurons were cultured in µ-Dish 35 mm dishes (Ibidi, 81156 or 81166 depending on the application) in Hybernate E (Gibco A1247601), 1× B27 Plus Supplement (ThermoFisher Scientific A3582801), 10 ng/ml BDNF (PeproTech 450-02), 10 ng/ml NT-3 (PeproTech 450-03), 1 mg/ml mouse laminin (Invitrogen 23017015), and 2 μg/ml doxycycline.

### Mitochondrial fission after axotomy

iPSCs were differentiated as described above. On day 1 of differentiation, cells were transduced with the appropriate lentivirus. Neuronal induction media was changed 24 hours later. On day 3 of differentiation, 50,000 cells were plated in a 5 µL drop of neuronal maturation media in the middle of a 35mm dish (ibidi, 81156) for the axons to grow out from the center. Dishes were incubated for 20 minutes at 37°C to allow cells to attach, and 2mL of neuronal maturation media was then added to each well. Cells were fed with half-media changes every two days until the day of the experiment. Laser axotomies were performed as described above. Single axons were randomly selected. Five images were taken before axotomy every 5 seconds and axons were imaged for 5 minutes after axotomy every 5 seconds. The number of mitochondrial particles was quantified using imageJ imaging software and normalized to the number of mitochondria pre-axotomy. When treating neurons with different chemical compounds, neurons were pretreated for one hour prior to performing axotomies. The following compounds were used GNE3511 (500 nM, Sigma, 5331680001), JNKi (SP600125 1µM, Selleckchem S1460), P110 (1 µM, Tocris, 6897), BAPTA-AM (10mM Tocris 2787), BSTA1 (BAX activator, 10 µM, Sigma SML2243) and BAI1 (BAX inhibitor, 10 µM, Medchemexpress HY-103269).

### Axon separation and axotomies

For axon separation experiments after axotomy, 1.5 million cells were plated in 150 µl of maturation media in a strip in the center of a 6 cm dish. Cells were allowed to attach for 15 minutes at 37°C, after which 8 ml of media was added to the dish. Neurons were cultured for 1 month to allow for the axons to project outwards. Axotomy was performed using a sharp blade. Axons and cell bodies were separated and processed for western blotting.

### Immunostaining of cells

Cells were fixed in cold 4% PFA for 5-10 minutes, permeabilized using 0.1% TritonX in PBS for 5 mins and blocked in 5% normal donkey serum (NDS) in PBS at room temperature for 1 hour. Cells were immunostained with the selected primary antibody in 2.5% NDS overnight at 4°C. Cells were washed three times with PBS and stained with the appropriate secondary antibody at a concentration of 1:500 in 2.5% NDS for 1 hour at room temperature. Cells were washed three times with PBS and DAPI was used as a nuclear counterstain. Antibodies used against: βIII tubulin (Thermo Fischer, 2G10, 1:500), pDRP1 S637 (CST, 4867S, 1:1000), DLK (DLK antibody developed and kindly provided by Cell Signaling Technology, clone GF-23-1, 1:500), TOM20 (Santa Cruz, SC-17764, 1:500).

### Calcium imaging

Neurons were plated as described above in the center of a 35mm dish, and their axons were allowed to grow outwards. 10-14 days post induction, neurons were treated with Fluo-4 AM cell permeable calcium dye (ThermoFischer, F14201) following the manufacturers indications. Axotomies were performed on single axons as described above and imaged every 2 seconds for 2 minutes after axotomy. Fluorescence intensities were calculated using ImageJ software and normalized to pre-axotomy levels.

### HEK cell culture and transfection

HEK-293T cells were cultured in DMEM (Gibco, cat. no 11995065) supplemented with 10% FBS (Gibco; 10437028). For transfection, cells were dissociated using Trypsin-EDTA (0.25%) (ThermoFischer, 25200056) and plated in DMEM, 10%FBS. The day after plating cells were transfected with 1 µg DNA plasmid, 8 µl lipofectamine 2000 (Invitrogen, 11668030) for 24 hours. For western blotting, cells were harvested in ice cold RIPA buffer (Invitrogen, 89901) supplemented with protease inhibitors (Sigma, 11836153001) and PhosStop (Sigma 4906845001). For immunostaining, cells were fixed in 4% PFA for 10 minutes.

### Western blotting

Protein concentrations were calculated using a BCA assay (ThermoFisher 23225). 10-20 µg of total protein lysate was loaded into a 4–20% Mini-PROTEAN® TGX™ Precast Protein Gels (Biorad 4561095). Gels were transferred into PVDF membranes and blocked for 1 hour at room temperature in 5% BSA. Membranes were incubated with the appropriate primary antibody in 2.5% BSA solution in a cold room, rocking, overnight. Membranes were washed 3 times with PBS 0.1% Tween solution. Secondary antibody was incubated for 1 hour at room temperature at a concentration of 1:8000 in 2.5% BSA solution. Membranes were washed 3 times with PBS 0.1% Tween solution prior to developing. Western blots were developed using Clarity Western ECL Substrate (Biorad 1705061) and imaged using a ChemiDoc MP Imaging system (Biorad). Band intensity was quantified using FIJI imaging analysis software. The following antibodies were used at the concentrations stated: anti-DRP1 (Santa Cruz Biotechnology, sc-101270, 1:1000), anti-pDRP1 S616 (CST 3455S, 1:1000), anti-ß Actin (Sigma A1978, 1:5000), anti-p-cJun S63 (CST 9261, 1:1000), anti-pDRP1 S637 (CST 4867S, 1:1000).

Abundance of phospho-DRP1/ total DRP1 was obtained running two western blots loaded with the same amount of total protein. Phosphorylated and unphosphorylated DRP1 protein bands were each normalized to actin levels, then the value for pDRP1 was divided by the value for DRP1 to obtain the relative levels of phosphorylated protein.

### Axon Degeneration index and neuron survival

Induction of iPSCs into neurons was performed as described above. On day 1 of differentiation, neurons were transduced with a cytoplasmic mApple (cyto mApple)-expressing lentivirus, and the media was changed 24 hours later. On day 3 of differentiation, transduced neurons were mixed with mixed with untransduced neurons at a ratio of 1:8000 to achieve sparse labelling of neurons, and re-plated on a 35mm gridded dish (Ibidi 81166). On day 14 of differentiation, the positions of selected axons or neuron somas was recorded and laser axotomies were performed as described above. Dished were stored in a temperature and CO_2_ controlled incubator until imaged next. Injured axons and somas were re-located at different time points after axotomy using the gridded dish and imaged. Axons were injured <1mm away from the soma. The axon degeneration index was calculated as previously described [37]. The following compounds were used: JNKi (SP600125 1µM, Selleckchem S1460), Actinomycin D (ActD) (Selleckchem S8964; 3µM).

### Airyscan imaging

Neurons were plated in 1mm German Glass Coverslips thickness #1.5 (Electron Microscopy Sciences 72290-04). 10-day old neurons were fixed with 4% PFA for 10 minutes at room temperature. Cells were permeabilized with 0.1% Triton-X for 5 minutes and blocked in 5% normal donkey serum (NDS) at room temperature for an hour. Primary antibody incubation was carried out in 2.5% NDS overnight at 4°C. Cells were washed three times with PBS and stained with the appropriate secondary antibody at a concentration of 1:500 in 2.5% NDS for 1 hour at room temperature. Cells were washed three times with PBS and DAPI was used as a nuclear counterstain. Coverslips were mounted using ProLong Glass antifade (Invitrogen P36980).

### Seahorse analysis

For Seahorse analysis, 50,000 cells per well were plated after 3 days of induction in a specialized Seahorse analyzer 96-well plate (Agilent, 103774-100). Cells were washed in PBS once and incubated in a hypoxic chamber following the manufacturer’s procedure. Seahorse analysis was performed using a Seahorse XFe 96 Analyzer (Agilent). The average value for three wells was used for analysis.

### Protein purification and *in vitro* kinase assay

DLK and DRP1 were cloned into a pcDNA3-C-terminal GST vector. 12 million HEK293 cells were plated in 10 cm dished coated with Matrigel. The day after, cells were transfected with 10 µg plasmids and 40 µl lipofectamine. 24 hours post transfection, cells were washed twice with PBS and harvested in 880 µl ice cold RIPA buffer (Invitrogen, 89901) supplemented with protease inhibitors (Sigma, 11836153001) and PhosStop (Sigma 4906845001). Lysates were sonicated and spun at 18000 g 10 minutes at 4°C and the pellet was discarded. 500 µl Glutathione-Sepharose beads (Sigma GE17-0756-01) were washed twice in 1ml in lysis buffer. Washed beads to 1 ml lysate and incubate end-over-end shaking in cold room overnight. The next morning. Beads were washed 4 times with lysis buffer + 0.4M NaCl. The protein-containing beads were pelleted by spinning 1min 8000 rpm at 4°C, the supernatant was discarded, and the beads were resuspended in fresh 200 mM glutathione 1X PBS solution for elution of the desired protein. The beds were incubated for 20 minutes at room temperature shaking. The eluted protein and the beads were separated by briefly spinning in a SpinX column tubes (Sigma CLS8162-24EA). The purity of purified protein was by running 5 µg of protein on an SDS PAGE gel and performing Comassie stain. The purified protein was stored at -80°C. For *in vitro* kinase assay, purified GST-DLK or GST-DLK-S302A was mixed with purified GST-DRP1 at a 1:1 ratio in 50 µl kinase buffer (50mM TrisHCL pH 7.5, 150mM NaCl, 10mM MgCl, 10mM MnCl_2_, 1.8mM ATP) incubated for 60 mins at 37°C. In vitro kinase assay was also performed in the presence of DLKi inhibitor GNE3511 (500 nM, Sigma, 5331680001) and JNKi (SP600125 1µM, Selleckchem, S1460). The reaction was terminated by adding SDS gel loading buffer and the sample was subjected to western blotting.

### Phospho-mass spectrometry

DRP1 bands, with ∼0.5 µg protein on each band, were cut from SDS-PAGE. In-gel samples were reduced with 10 mM Tris(2-carboxyethyl)phosphine hydrochloride, alkylated with N-Ethylmaleimide, and digested with trypsin at 37°C for 18 hours. Peptides were extracted, desalted and injected into a nano-LC-MS/MS system where an Ultimate 3000 HPLC was coupled to an Orbitrap Lumos mass spectrometer (Thermo Scientific) via an Easy-Spray ion source (Thermo Scientific). Peptides were separated on an ES902 Easy-Spray column (75-μm inner diameter, 25 cm length, 3 μm C18 beads; Thermo Scientific) with mobile phase B (0.1% formic acid in LC-MS grade acetonitrile) increased from 3% to 24% in 60 min. The flow rate was maintained at 300 nl/min. MS and MS/MS data were acquired on Thermo Scientific Orbitrap Lumos mass spectrometer. MS1 scans were acquired in orbitrap at of a resolution 120k with a mass range of m/z 400-1500. MS2 scans were acquired in ion trap with ETciD method at normal scan rate. The isolation width was 1.6 m/z. Ions were excluded after 1 acquisition and the exclusion duration is 9 sec. MS1 scans were performed every 3 sec. As many MS2 scans were acquired within each MS1 scan cycle. Proteome Discoverer software version 2.4 was used for protein identification and quantification. Database search was performed against Sprot Human database using Mascot search engine. Mass tolerances for MS1 and MS2 scans were set to 5 ppm and 0.4 Da, respectively. Up to 2 missed cleavage was allowed for trypsin digestion. NEM on cysteines was set as fixed modification. Oxidation (M) and Phosphorylation (STY) were searched as variable modifications. Spectra of phosphopeptides matched by database search were manually checked. The search results were filtered by a false discovery rate of 1% at the protein level. Proteins detected with 1-2 peptides were further filtered out. Protein abundance values were calculated by summing the abundance of unique and razor peptides matched to that protein. Each sample group contains 4 replicates. The ratios were calculated using protein abundance without normalization.

### Electron microscopy

Neurons were plated in 1mm German Glass Coverslips thickness #1.5 (Electron Microscopy Sciences, 72290-04). 10 Day old neurons were fixed with in 4% PFA, 1% Glutaraldehyde for 40 minutes. Fixed cells were placed in phosphate buffer containing 0.1% sodium borohydride to inactivate residual aldehyde groups. Cells were then washed with phosphate buffer several times until the solution was clear of bubbles. To improve reagent penetration, the cells were then treated with phosphate buffer containing 0.05% Triton X-100. To prevent nonspecific binding of the immunoreagents, cells were incubated in Aurion blocking solution for Goat gold conjugates (Electron Microscopy Sciences, Hatfield, PA., containing phosphate buffer, pH 7.4, normal goat serum (NGS), bovine serum albumin (BSA), and cold water fish skin gelatin (CWFG)). After blocking, cells were incubated in the primary antibody, DLK (custom antibody from Cell Signaling Technology, clone GF-32-1, 1:100) diluted with incubation buffer (phosphate buffer containing 0.2% Aurion acetylated bovine serum albumin (BSA-c), Electron Microscopy Sciences, Hatfield, PA., 5% CWFG and 5% NGS, pH 7.4) for two hours at room temperature. After washes with incubation buffer, cells were incubated in the secondary antibody Aurion ultra-small gold-conjugated F(abʹ)_2_ fragments of Goat anti-Rabbit IgG, (Electron Microscopy Sciences, Hatfield, PA.) diluted 1:100 with incubation buffer overnight at 4°C. To remove unbound secondary antibody, cells were washed thoroughly with incubation buffer and then with PBS. After washes, cells were prepared for silver enhancement.

### Silver Enhancement and Cell processing for Electron Microscopy

Cells were washed with MilliQ water and then transferred to Aurion R-Gent SE-EM silver enhancement solution, Electron Microscopy Sciences, Hatfield, PA. and incubated at room temperature for 35 minutes. The enhancement was stopped by washing cells several times in MilliQ water. The following processing steps were carried out using the variable wattage Pelco BioWave Pro microwave oven (Ted Pella, Inc., Redding, CA.): cells were rinsed in MilliQ water, post-fixed in 1% osmium tetroxide made in MilliQ water, ethanol dehydration series up to 100% ethanol, followed by a Embed-812 resin (Electron Microscopy Sciences, Hatfield, PA.) infiltration series up to 100% resin. The epoxy resin was polymerized for 20 hours in an oven set at 60°C. Ultra-thin sections (90nm) were prepared on a Leica EM UC7 ultramicrotome. Ultra-thin sections were picked up and placed on 200-mesh cooper grids (Electron Microscopy Sciences, Hatfield, PA) and post-stained with uranyl acetate and lead citrate. Imaging was performed on a JEOL-1400 Transmission Electron Microscope operating at 80kV and images were acquired on an AMT BioSprint 29 camera.

### Generation of DLK KO iPSCs

WT iPSCs were transfected with Cas9-GFP, and two gRNA containing plasmids targeting exons 3 and 5. gRNA expressing plasmids were purchased from Sigma Aldrich (U6-gRNA:hPGK-puro-2A-tBFP, gRNA1: CCCAGGCTCCCTGCTACTGCAT, gRNA 2: TCCTTTGGCGTGGTGCTATGGG). 1 day after transfection, expression of Cas9 and gRNAs was validated by the expression of GFP and BFP respectively and iPSCs were treated for 3 days with 10 µM Puromycin (Invitrogen A11113803) and allowed to recover for 3 days. Cells were then dissociated with accutase for 10 minutes and dissociated my pipetting forcefully 10 times. A 1:2 serial dilution was then performed in two 6 well plates. iPSCs were fed every 3 days until individual colonies were figure visible (usually 10-14 days). Individually colonies were picked using a P1000 pipette and transferred into individual wells in a 24-well plate. Individual clones were allowed to expand, passaged and their DNA was extracted for DLK KO validation using PCR (Forward primer: TCAGGTGAATGCTGAGCCAGCT, reverse primer: TGGAGACTGTTGCTTCCCACAC). A shorter band from ∼1200bp to ∼200bp was expected if the KO was successful. Potential clones were sequenced. Further validation was performed by differentiating potential clones into neurons and knock out of DLK was validated by western blotting and immunofluorescence.

### Animals

All animal care and experimental procedures were performed in accordance with animal study proposals approved by the National Institute of Child Health and Human Disease Animal Care and Use Committee (animal protocol number 20-003, Le Pichon lab), National Eye Institute (animal protocol number NEI-606, Li lab) and by the Baylor College of Medicine ACUC (animal protocol #AN-7208; Watkins lab). C57Bl/6J (Jax#000664) mice of both sexes, aged between 6-10 weeks old, were used in this study.

### Mouse optic nerve preparation for Western blotting

Mice were anesthetized with 2.5% avertin followed by decapitation. Each eye was carefully removed from orbit with the optic nerve attached. The proximal segment of the injured nerve was harvested and dissected on a cold (4°C) surface in PBS containing protease inhibitors (Sigma 11836153001) and PhosStop (Sigma 4906845001). Nerves from 4-6 mice were pooled to obtain enough protein for each lane of a Western blot. Nerves were mechanically lysed in 40 µl ice cold RIPA buffer (Invitrogen 89901) supplemented with protease inhibitors (Sigma 11836153001) and PhosStop (Sigma 4906845001) for 2 minutes using a micro pestle. Nerves were then sonicated for 15 minutes at 4°C. Protein lysates were centrifuged at 18 000g, 4°C for 10 minutes and the pellet was discarded. The samples were subjected to the western blotting procedure described above.

### Adeno-associated virus (AAV)

Adeno-associated viral vectors carrying human synapsin-1 (*hSyn1*) promoter driving expression of saCas9-U6-control, saCas9-U6-DRP1-gRNA, or eGFP were generated by Epoch Life Sciences Inc. The plasmids were then packaged into AAV2 capsids at the Optogenetics and Viral Vector Core at Duncan Neurological Research Institute, Houston.

### Intravitreal injections

Mice to be injected were anesthetized with 3% isoflurane. The eyes were sterilized by 3 repeated applications of 5% ophthalmic betadine (Henry Schein #6900250), Opti-clear ophthalmic eye wash (Akorn #NDC 17478-620-04) and a dry wipe. A topical anesthetic 0.5% proparacaine HCl ophthalmic solution (Henry Schein #1365345) was applied. The eyeball was punctured using a 5-µl Hamilton syringe loaded with a custom 33-gauge needle (Hamilton #7803-5) and some of the intraocular pressure was relieved. The same puncture site was used to reinsert the Hamilton needle and 2 µl of AAV in titers ranging from 10^12^ -10^13^ vg/ml was delivered per eye.

### Intra-orbital optic nerve crush (ONC)

Two weeks after intravitreal injections, animals underwent optic nerve crush surgeries. Mice undergoing surgery were dosed with 1mg/kg buprenorphine sustained release formulation 1 hour before surgery. Right before surgery, mice were anesthetized with 3% isoflurane. The non-surgical eye received artificial tears ointment (Covetrus #11695-6832-1. The surgical eye was sterilized as described in the intravitreal injections section above. Topical anesthetic 0.5% proparacaine HCl ophthalmic solution (Henry Schein #1365345) was applied on the surgical eye (left). A pair of Vannas scissors (World Precision Instruments #501777) were used to make incisions in the conjunctival layers. The optic nerve section in the intra-orbital space was exposed by using two pairs of suture-tying forceps (Fine Science Tools #1106307) to gently clear the soft tissue behind the eye until the optic nerve was visible. The optic nerve was manually crushed for 5s by using a pair of Dumont forceps (Fine Science Tools #1125325). The eyeball was gently placed back into the orbit. Animals were post-surgically monitored in their home cages until sternal recumbency was observed.

### Immunolabeling of retinae for cleaved caspase 3

At the experimental endpoint (3 days after optic nerve crush) mice were euthanized by an overdose of isoflurane and followed by decapitation. The eyes were enucleated and drop-fixed in 4% paraformaldehyde for 1 hour. The retinae were dissected out in 1X phosphate-buffered saline (PBS) and then blocked with blocking buffer (5% goat serum, 0.5% Triton X-100 and 0.025% sodium azide in 1X PBS) for 30min. The retinae were then immersed in primary antibody solution (anti-cleaved caspase 3, CST 9661) prepared by dilution in blocking buffer solution and incubated for 5 days in 4°C. They were then washed three times in 1X PBS with 0.5% TritonX-100 for 30 min each, and then moved to appropriate secondary antibody solution prepared in blocking buffer and incubated overnight in 4°C. After secondary antibody incubation, they were washed again three times in 1X PBS with 0.5% TritonX-100 for 30 min each. Retinae were then mounted in Drop-n-Stain EverBrite^TM^ Mounting Medium (Biotium #23008) onto slides and imaged using Zeiss Axio Imager Z1 fluorescence microscope. 5-8 images were taken per animal in a blinded manner for quantification.

### Quantification of cleaved caspase 3-positive RGCs

The number of cleaved caspase 3 positive cells were quantified in 5-8 images per retina per animal in a blinded manner. The number of caspase 3 positive cells per section was averaged over all the images quantified to obtain the average number of cleaved caspase 3 cells per section.

### Retinal dissection for RBPMS analysis

For RGC survival 7 days post injury, prior to fixation in 4% paraformaldehyde for 1 hour, a burn signal was applied to the dorsal pole of enucleated eyeballs to maintain orientation. Retinas were then carefully dissected as flat whole-mounts employing four radial cuts, with the deepest incision previously marked at the dorsal pole. The flattened whole-mounts underwent an additional hour of post-fixation in 4% PFA, followed by a gentle removal of vitreous using brushes. Subsequently, they were kept in phosphate-buffered saline (PBS) for further processing.

### Immunofluorescence for RBPMS

Retinal ganglion cells (RGCs) in whole-mount retinas were identified through immunodetection of the RBPMS (RNA-binding protein with multiple splicing) protein. Initially, all retinas were permeated (4 × 10 minutes) in PBS containing 0.5% Triton X-100 (Tx). The primary antibody (rabbit anti-RBPMS, GeneTex #GTX118619) was diluted at 1:500 and incubated overnight with shaking at room temperature in a blocking buffer (2% Normal Donkey Serum, 0.5% Tx in PBS). Next, the retinas were washed and incubated overnight with the appropriate secondary antibody (donkey anti-rabbit Alexa 649, Jackson ImmunoResearch, #706-495-148) diluted 1:500. Finally, retinas were washed in PBS and cover-slipped vitreal side up with antifading mounting medium.

### Image acquisition

RBPMS^+^RGCs were imaged from flattened retinal whole-mounts using a 20x objective on an LSM 780 Zeiss confocal microscope equipped with a computer-driven motorized stage controlled by Zen Lite software (Black edition, Zeiss). Photomontage frames were captured contiguously side-by-side with an 8% overlap between them, and images within the same frame were acquired at intervals of 3.5 microns in the Z-dimension. The maximum-projection images were employed for subsequent analysis.

### Quantification of RBPMS RGCs

Quantification of RBPMS^+^RGCs was conducted on entire retinas using an automated algorithm in ImageJ. Briefly, to minimize interference with background labeling, a rolling ball radius of 50 pixels was subtracted. The application of a median radius of 1 smoothed the edges, followed by a maximum radius of 0.0005 to fill and dilate the objects. Subsequently, all images underwent an 8-bit grayscale transformation to eliminate color information, and a predetermined lower threshold generate a binary mask-like images. The “watershed” segmentation automatically separated particles that were in contact, while the “despeckle” median filter efficiently removed noise. Positive objects were counted within defined parameters regarding shape and size, to exclude those considered either too small or too large to be classified as RGCs. The automatic routine extracted cell counts and their corresponding coordinates (x, y), which were exported to a spreadsheet (Microsoft Office Excel; Microsoft Corp., Redmond, WA, USA) for further analysis (see the next section).

### Spatial Analysis

Topographical distributions of RBPMS^+^RGC densities were assessed using color-filled contour maps generated with Sigma Plot 13.0 for Windows (Systat Software, Inc., Richmond, CA, USA). These heat maps assigned a color code to each area of interest (83.3 × 83.3 μm) based on its density values, ranging from 0 (purple) to 3,600 RGCs/mm² (red) in a 10-step color scale.

### DLK inhibitor treatment

GNE-3511 solution was synthesized as previously described [71]. Briefly, GNE-3511 was dissolved in a solution of 0.5% w/v USP Grade Methyl Cellulose and 0.2% v/v Tween 80 in water up to 48 hours before dosing. Mice were dosed by oral gavage every 12 hours at 75 mg/kg with 10 ml/kg body weight of a 7.5 mg/ml solution of GNE-3511 or vehicle alone.

### Statistical analyses

All quantifications in this study (except for Western blots), were performed blind to experimental groups and conditions. All statistical analysis was performed in GraphPad Prism 9 unless otherwise stated.

## Acknowledgements

We would like to thank Drs. Vincent Schram and Chip Dye and the NICHD Microscopy core, Dr. Yan Li and the NINDS proteomics core, the NHLBI FACS core facility staff, Dr. Gareth Thomas (Temple University), Dr. Ana J. Garcia-Saez (Max Plank Institute of Biophysics) and Dr. Richard Cho (Cell Signaling Technologies) for reagents. We would also like to thank members of Dr. Michael Ward’s lab for advice, and Dr. Nicholas Ryba, Dr. Richard Youle, Dr. David Simon, Dr. Alexander Chesler, and Dr. Joseph Lewcock for helpful discussion and comments on the manuscript.

## Author contributions

Conceptualization: JGD and CLP. Methodology and Investigation: all authors. Formal Analysis: MN, FNN and JGD. Visualization: JGD and CLP.

Writing: JGD and CLP. Supervision: CLP, TW, LW, MEW.

Funding acquisition: CLP, TW, LW, MEW.

