## Supplemental for "DLK-dependent axonal mitochondrial fission drives degeneration following axotomy"

Supplementary Figure 1: Axotomy leads to calcium flooding of the axon and mitochondrial fission is Ca dependent.

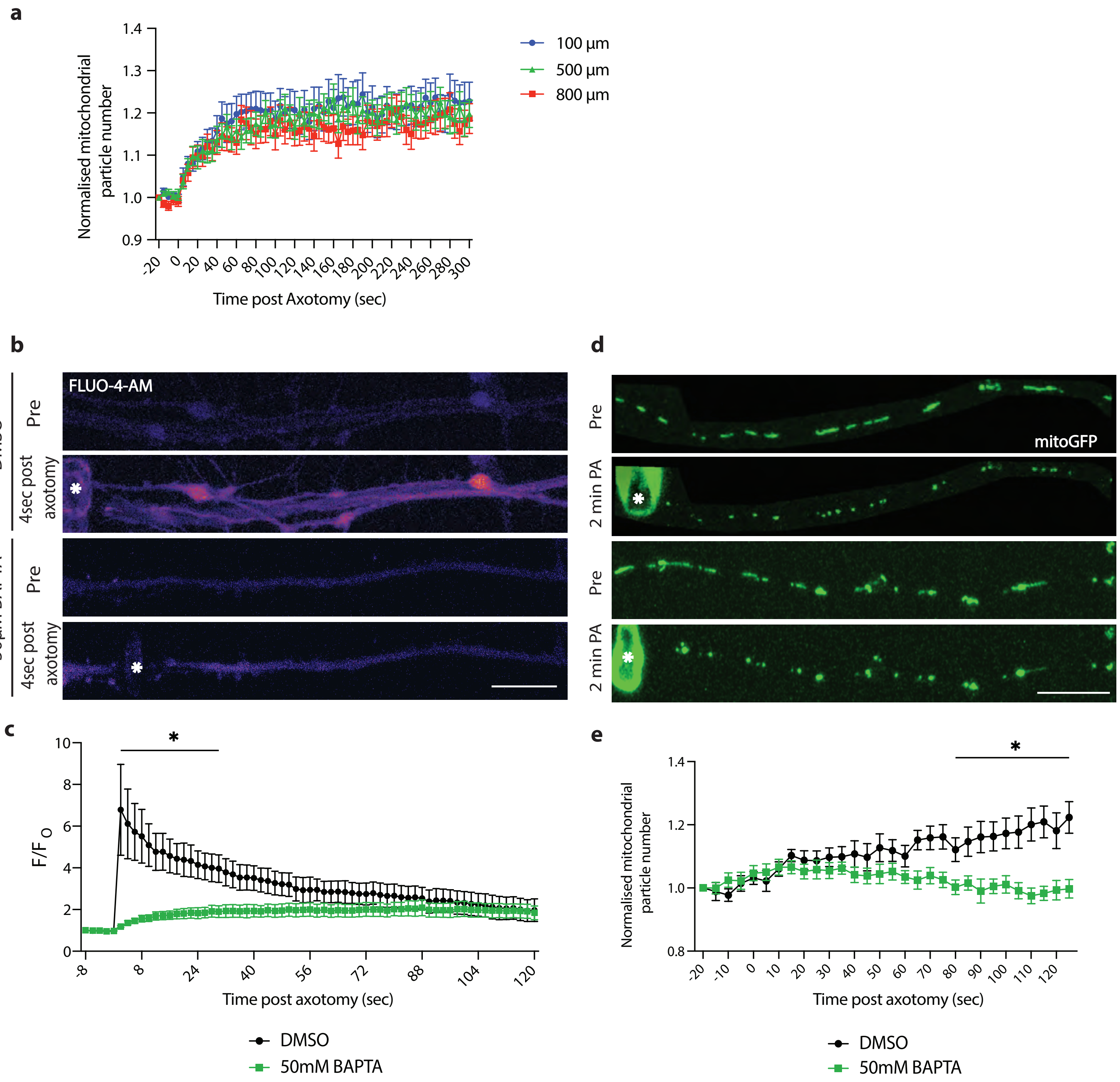

**Sup Figure 1 – Axotomy leads to calcium flooding of the axon and mitochondrial fission is Ca dependent.**

**a)** Normalized number of mitochondrial particles post axotomy in WT neurons axotomized 100µm (blue circle), 500µm (green triangle) and 800µm (red squares) from the soma. Results are represented as mean ± SEM. N=3 independent differentiations, N≥ 30 axotomized neurons.

**b)** Representative images of Fluo-4-AM calcium imaging of injured axons pre and 4 seconds post axotomy treated with DMSO and 50 µM BAPTA-AM. * Marks the site of axotomy. (Scale bar = 25 µm).

**c)** Relative Fluo-4-AM fluorescence after axotomy in DMSO and BAPTA AM-treated neurons after injury. Results are represented as mean ± SEM. N=6 cells. Two-way ANOVA, Bonferroni correction (p≤0.05 *).

**d)** Representative images of WT neurons transduced with mitoGFP (green) treated with DMSO and BAPTA-AM pre and 2 mins post axotomy (PA). * Marks the site of axotomy. (Scale bar = 25 µm).

**e)** Normalized number of mitochondrial particles post axotomy in DMSO (black) and BAPTA-AM (green)-treated neurons. Results are represented as mean ± SEM. N=2 independent differentiations, N≥ 16 axotomized neurons. (Two-way ANOVA, Bonferroni correction (p≤0.05 *)

Supplementary Figure 2: DLK localises to mitochondria in i3neurons

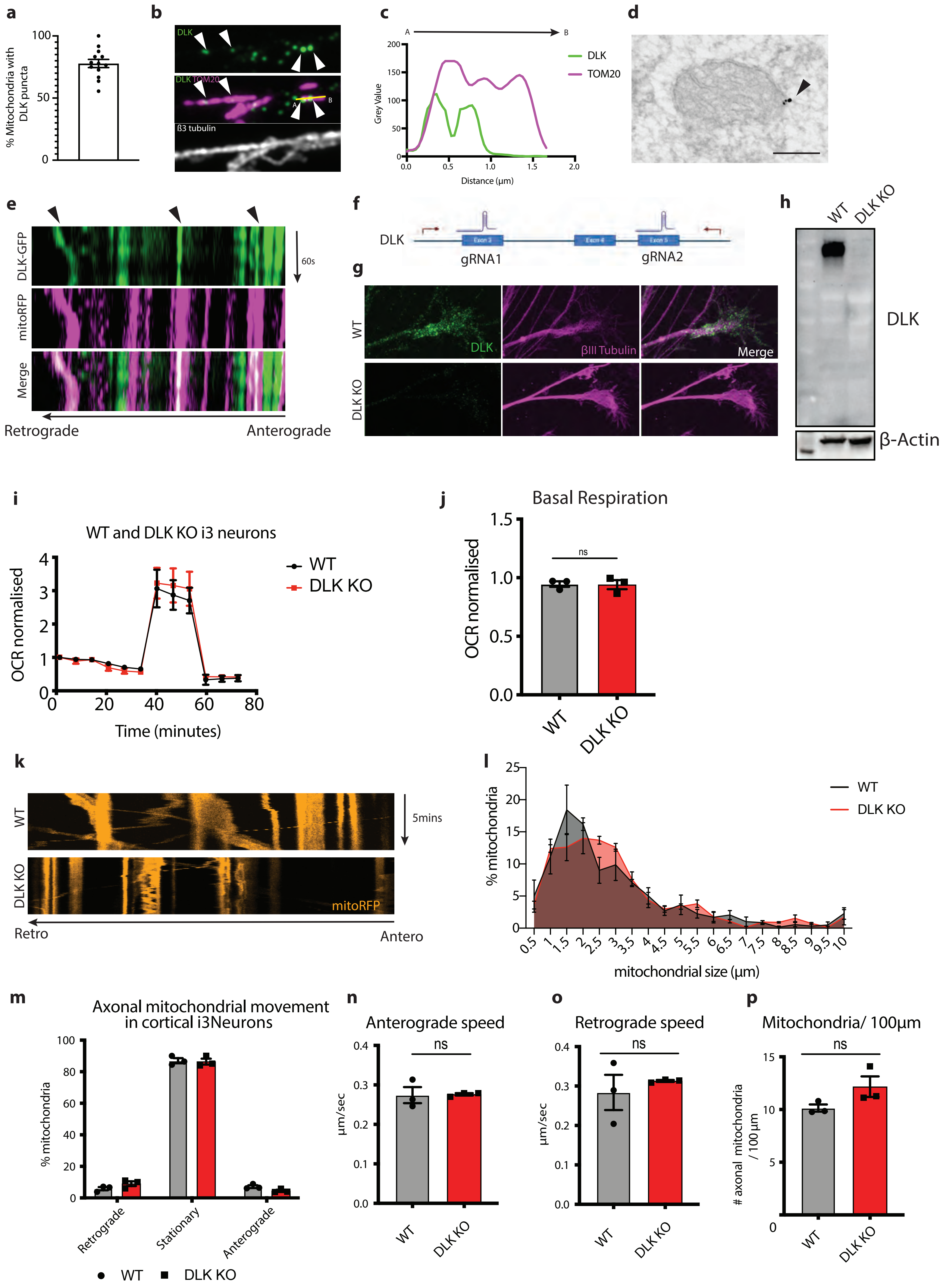

**Sup Figure 2 – DLK localizes to mitochondria in i3Neurons**

**a)** Quantification of percentage of axonal mitochondria containing DLK puncta. Results are represented as mean ± SEM. N=13 axons from 3 individual differentiations.
**b)** Representative high-resolution images of WT neurons axons stained with anti-DLK (green) and anti-TOM20 (magenta).
**c)** Line-scan analysis of relative fluorescence intensity from the dashed line shown in b.
**d)** Immuno-EM of endogenous DLK localized in the mitochondria of i3Neurons. Arrow indicates DLK puncta. (Scale bar = 5 µm).
**e)** Representative kymographs of DLK-GFP (green) and mitoRFP (magenta) co-trafficking along the axon.
**f)** Schematic representation of DLK knockout (KO) generation strategy. WT $i^3$ iPSCs were transfected with two gRNAs targeting DLK exon 3 and 5. Primers used for knockout validation are shown in red.
**g)** Representative images of WT and DLK KO neurons stained with DLK (green), ßIII tubulin (magenta).
**h)** Western blots of WT and DLK KO neurons. Immunoblot for DLK and loading control ß actin.
**i)** Seahorse oxygen consumption rate (OCR) analysis of WT and DLK KO i3Neurons normalized to WT Neurons. Results are represented as mean ± SEM. N=3 independent differentiations, 3 wells measured per condition.
**j)** Basal OCR levels of WT and DLK KO neurons normalized to UT $i^3$Neurons. Results are represented as mean ± SEM. N=3 independent differentiations, 3 wells measured per condition. No significant changes observed (ns).
**k)** Representative kymographs of WT and DLK KO axon mitochondrial (mitoRFP orange) movement.
**l)** Quantification of WT and DLK KO axon mitochondrial length. Results are represented as mean ± SEM. N=3 independent differentiations, N≥30 axons. No significant differences observed (ns).
**m)** Quantification of percentage of stationary, retrograde and anterograde-moving mitochondria in WT and DLK KO neurons. N=3 independent differentiations, N≥ 30 axons. No significant differences observed (ns).
**n)** Quantification of anterograde mitochondrial speed in WT and DLK KO neurons. N=3 independent differentiations, N≥ 30 axons. No significant differences observed (ns).
**o)** Quantification of retrograde mitochondrial speed in WT and DLK KO neurons. No significant differences observed. N=3 independent differentiations, N≥ 30 axons. No significant differences observed (ns).
**p)** Quantification of the number of mitochondria per 100 µm in WT and DLK KO axons. No significant differences observed. N=3 independent differentiations, N≥ 30 axons. No significant differences observed (ns).

Supplementary Figure 3

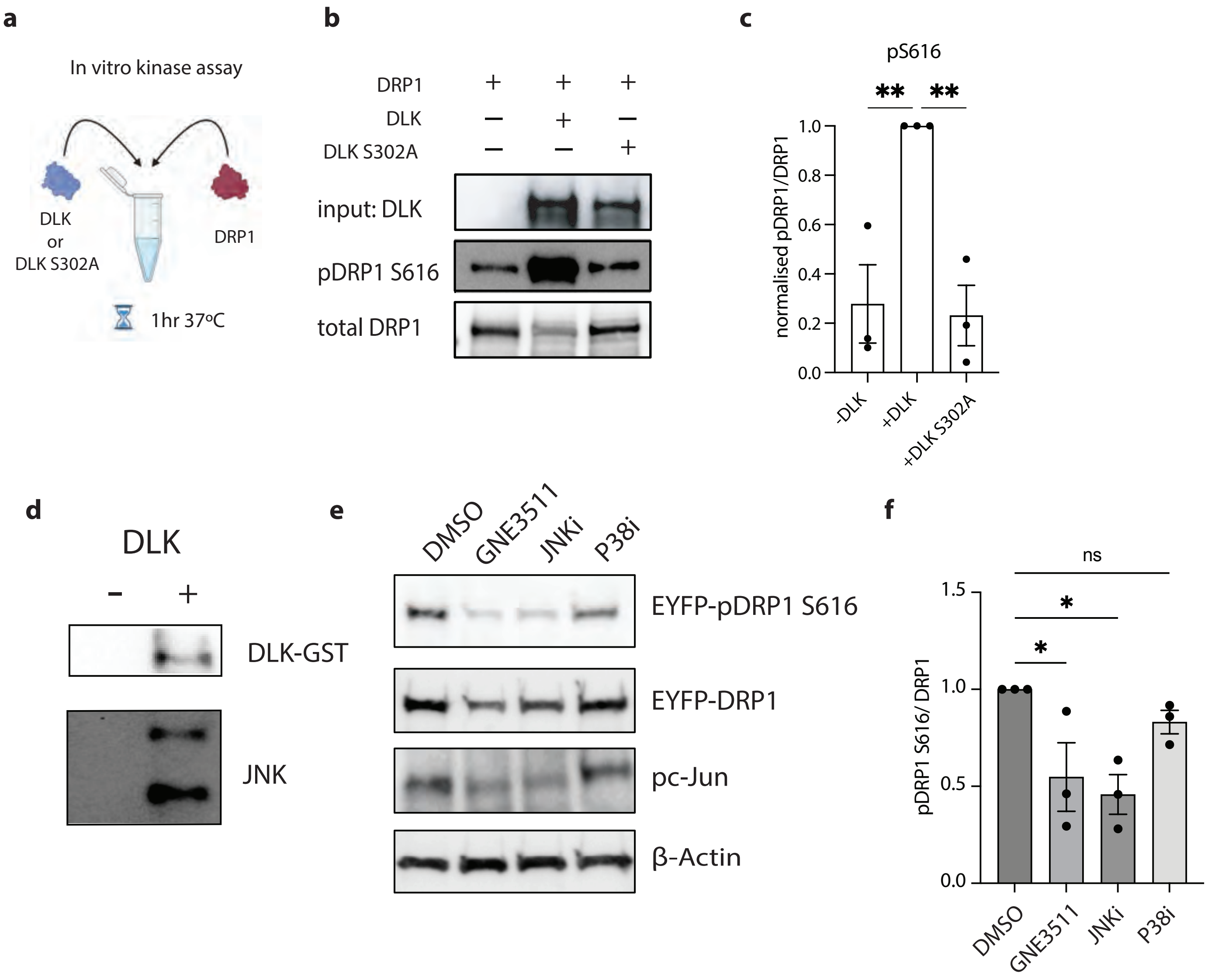

**Sup Figure 3 – the DLK/JNK pathway phosphorylates DRP1 in vitro**

a) Schematic representation of the *in vitro* kinase assay using purified DLK, DLK S302A and DRP1 to test the direct phosphorylation of DRP1 by DLK.
b) Representative western blots of *In vitro* kinase assay utilizing separately purified DRP1, WT DLK and kinase dead DLK S302A. Representative immunoblots for pS616-DRP1, total DRP1 and DLK
c) Quantification of Quantification of pS616-DRP1/total DRP1 in an *in vitro* kinase assay. Results are represented as mean ± SEM. One-way ANOVA, Bonferroni correction ($p \leq 0.01$ **).
d) Representative western blot of JNK co-precipitating with purified DLK-GST in HEK-293T cells. Immunoblots for total DLK and JNK.
e) Representative western blots of HEK293A cells transfected for 24 hours with DLK-GFP and EYFP-DRP1 and treated with DMSO, GNE-3511, JNKi and P38i. Immunoblot for pS616-DRP1, total DRP1, pS63-cJun and loading control ß actin.
f) Quantification of pS616-DRP1/total DRP1 levels after 24-hour expression of DLK-GFP and EYFP-DRP1 treated with DMSO, GNE-3511, JNKi and P38i. Results are represented as mean ± SEM. One-way ANOVA, Bonferroni correction (not significant (ns), $p \leq 0.05$ *).

Supplementary Figure 4

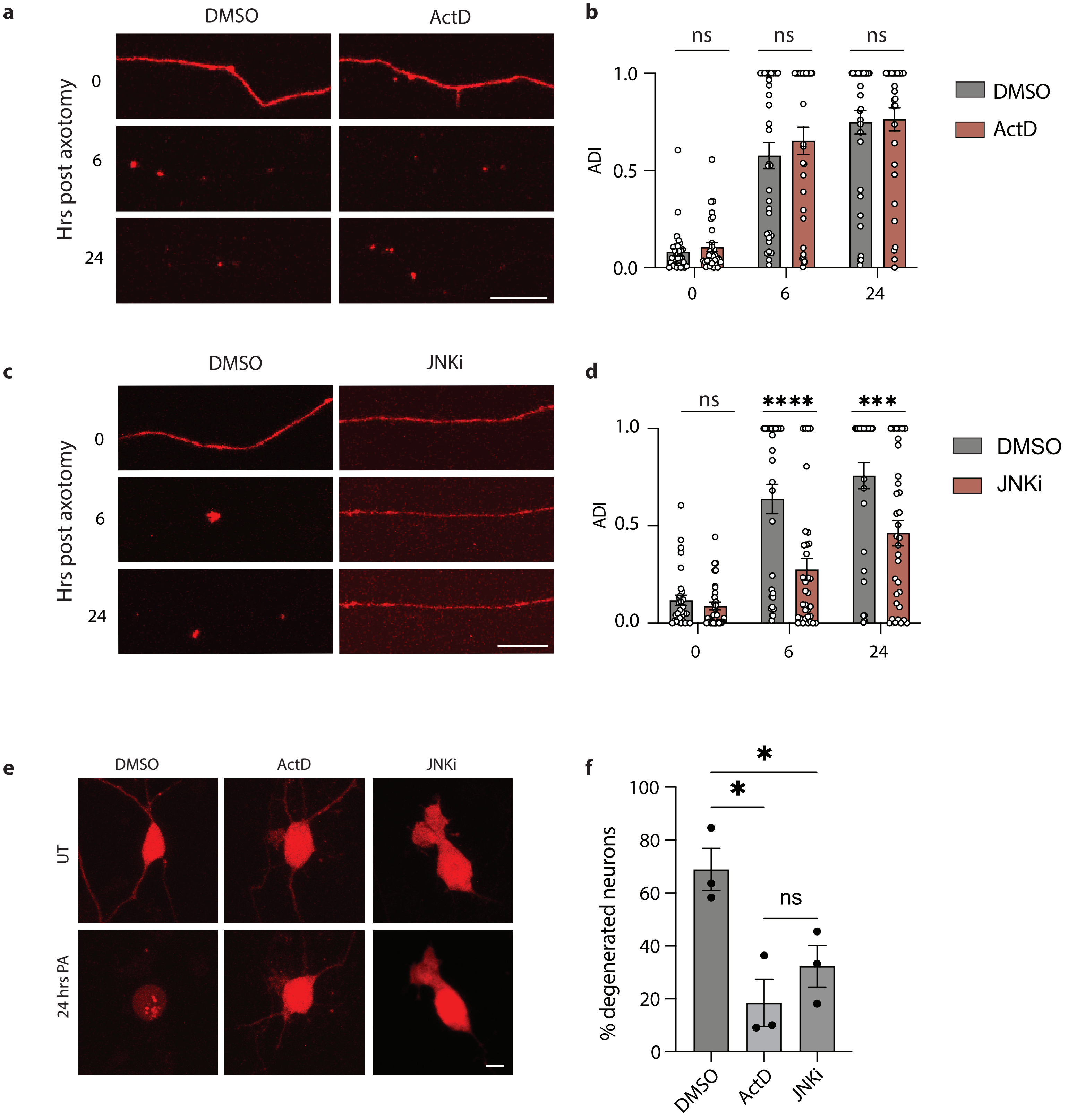

**Sup Figure 4 – Effect of blocking transcription and JNK on cell death and axon degeneration after axotomy**

a) Representative images of neuron axons treated with DMSO or Actinomycin D (ActD) transduced with cyto mApple (red) proximal to the site of injury 0, 6 and 24 hours post axotomy (Scale bar = 25 μm).
b) Quantification of axon degeneration index (ADI) in neurons treated with DMSO or Actinomycin D (ActD) 0, 4, 8 and 24 hours post axotomy. Results are represented as mean ± SEM. N=3 independent differentiations, N≥ 32 axons. Two-way ANOVA, Bonferroni correction (not significant, ns).
c) Representative images of neuron axons treated with DMSO or JNKi transduced with cyto mApple (red) proximal to the site of injury 0,4, 8 and 24 hours post axotomy (Scale bar = 25 μm).
d) Quantification of axon degeneration index (ADI) in neurons treated with DMSO or JNKi 0, 4, 8 and 24 hours post axotomy. Results are represented as mean ± SEM. N=3 independent differentiations, N≥ 31 axons. Two-way ANOVA, Bonferroni correction ($p \leq 0.005$ ***, $p \leq 0.001$ ****).
e) Representative images of neuron cell bodies transduced with cyto mApple (red) treated with DMSO, Actinomycin D (ActD) and JNKi pre and 24 hours post axotomy (PA). (Scale bar = 40μm).
f) Percentage degenerated neurons treated with DMSO, Actinomycin D (ActD) and JNKi 24 hours post axotomy. Results are represented as mean ± SEM. N=3 independent differentiations. Unpaired t-test (not significant, ns. $p \leq 0.05$ *).
